# Discovery and Characterization of a Chemical Probe for Cyclin-Dependent Kinase-Like 2

**DOI:** 10.1101/2024.05.12.593776

**Authors:** Frances M. Bashore, Sophia M. Min, Xiangrong Chen, Stefanie Howell, Caroline H. Rinderle, Gabriel Morel, Josie A. Silvaroli, Carrow I. Wells, Bruce A. Bunnell, David H. Drewry, Navjot S. Pabla, Sila K. Ultanir, Alex N. Bullock, Alison D. Axtman

## Abstract

Acylaminoindazole-based inhibitors of CDKL2 were identified via analyses of cell-free binding and selectivity data. Compound **9** was selected as a CDKL2 chemical probe based on its potent inhibition of CDKL2 enzymatic activity, engagement of CDKL2 in cells, and excellent kinome-wide selectivity, especially when used in cells. Compound **16** was designed as a negative control to be used alongside compound **9** in experiments to interrogate CDKL2-mediated biology. A solved co-crystal structure of compound **9** bound to CDKL2 highlighted key interactions it makes within its ATP-binding site. Inhibition of downstream phosphorylation of EB2, a CDKL2 substrate, in rat primary neurons provided evidence that engagement of CDKL2 by compound **9** in cells resulted in inhibition of its activity. When used at relevant concentrations, compound **9** does not impact the viability of rat primary neurons or certain breast cancer cells nor elicit consistent changes in the expression of proteins involved in epithelial–mesenchymal transition.

---

Cyclin-dependent kinase-like 2 (CDKL2, KKIAMRE, P56) is a serine/threonine kinase that belongs to the CMGC kinase group and CDKL family.^1^ The CDKL family consists of five members: CDKL1–5. Enhanced tissue expression of CDKL2 is observed in the retina and testis, but it is also non-specifically expressed throughout the brain and in the lungs and kidneys.^2-4^ This kinase is cytoplasmic and localizes to the nucleoplasm and centrosome in cells.^5-7^ Based on characterization of animal and human cDNA clones, at least four variants of this enzyme may exist, generated by alternative splicing.^2^ All putative isoforms identified from human, rabbit, and mouse cDNAs contain a common kinase domain and vary in the carboxy-termini regions.^2^ In humans, two major transcripts were found in the adult testis, kidney, brain, and lung, and a single transcript in the fetal brain and kidney.^7^

While few publications have been published that focus on CDKL2, some intriguing details about the biology of this kinase have been described. CDKL2 is activated in cells by treatment with epidermal growth factor.^7^ As this activation does not require phosphorylation of the conserved MAP kinase dual phosphorylation motif (Thr-Asp-Tyr) of CDKL2, CDKL2 is not considered a functional member of the MAP kinase group.^7^ While several studies have provided a link between CDKL2 and tumorigenesis and have highlighted that the function of CDKL2 in different cancer types is distinct, an understanding of how CDKL2 modulates oncogenic progression is not yet understood. Reports have linked CDKL2 expression to the progression and survival of patients with kidney^6, 8^, stomach^5, 9, 10^, liver^11^, brain^12^, prostate^13^, and breast^14-16^ cancers. CDKL2 expression is a favorable prognostic marker in glioma and renal and liver cancers.^6, 8, 11, 12^ Increased DNA methylation of CDKL2 has been observed in tissue taken from patients with liver^11, 17, 18^ and prostate^13^ cancer. In hepatocarcinomas, CDKL2 hypermethylation has been correlated with its reduced expression as well as progression of the cancer.^11^

However, in breast cancer, including invasive subtypes, high CDKL2 expression is associated with poorer survival.^5^ Based on an orthotopic breast cancer xenograft model, this is driven, in part, by CDKL2 promoting primary tumor formation and spontaneous metastasis.^5^ CDKL2 was identified as a regulator of epithelial–mesenchymal transition (EMT), a process associated with increased migration, metastasis, and therapeutic resistance and cells becoming stem cell-like.^5^ Accordingly, CDKL2 expression was significantly higher in mesenchymal breast cancer cell lines, human mesenchymal stem cells, and fibroblasts when compared to epithelial breast cancer cell lines.^5^ In contrast to these findings, it was reported that CDKL2 is hypermethylated and its expression downregulated in HER2+ breast cancer tissues. Accordingly, CDKL2 upregulation was suggested to inhibit cancer progression.^15^ Based on these results, the role of CDKL2 in breast cancer could be nuanced and/or more research needs to be done in this area. Similarly, based upon analysis of patient samples, the literature is mixed on the role of CDKL2 in gastric cancer, with one group reporting that high CDKL2 mRNA level predicts shorter overall survival^9^ and two other groups reporting that loss of CDKL2 predicts poor prognosis.^5, 10^ CDKL2 expression, therefore, may not be a reliable prognostic marker in some cancers.

Beyond cancer, CDKL2 plays a role in development, supporting behavior control, emotion, and cognitive functions required to acquire spatial and contextual learning.^4, 19^ CDKL2 is also responsive to viral infection. HSV-2 infected HeLa cells demonstrated reduced expression of CDKL2, which was suggested to impact processes such as apoptosis and cell cycle in these cells.^20^ With better tools, including a small molecule chemical probe, additional studies can be performed to further characterize how CDKL2 modulates other disease-relevant pathways.

From a structural standpoint, crystal structures of the CDKL2 kinase domain bound to TCS 2312 (PDB code: 4BBM) and to CDK1/2 Inhibitor III (PDB code: 4AAA) have been solved and deposited.^21^ These structures revealed that CDKL2 has a distinct C-terminal αJ helix that is also found in CDKL3 but not in other CDKL family members.^21^ The motif occludes the recruitment site for MAPK substrates, supporting the idea that CDKL family members mediate disparate protein interactions when compared to MAPKs.^21^ The αJ region was also found to be essential for CDKL2 function and its deletion significantly reduced CDKL2 activity. The ligand-bound structures harbored an inactive αC-out conformation and ligand binding was stabilized by a collapsed P-loop conformation.^21^ Interestingly, an active kinase conformation was also compatible with binding of CDK1/2 Inhibitor III and thus inhibitor interactions are not solely responsible for the inactive conformation observed.^21^ It is worth noting that structures of other CDKL family members displayed characteristics of active kinases, making CDKL2 structurally unique.^21^

Importantly, the compounds that were co-crystallized with CDKL2 are not selective for CDKL2, but rather broad-spectrum ATP-competitive kinase inhibitors. While these do not represent the only kinases inhibited by the respective compounds, TCS 2312 is sold as an inhibitor of CHK1, while CDK1/2 Inhibitor III is often used to inhibit CDK1 and CDK2 as its name implies. No potent and selective small molecule inhibitors of CDKL2 have been published. Such a compound would represent a powerful tool that would aid in deciphering the biological roles of CDKL2.

In search of a high-quality CDKL2 inhibitor, we reviewed our kinome-wide screening data. We identified the acylaminoindazoles as a promising chemical series with robust CDKL2 binding affinity. The acylaminoindazoles were the source of our AAK1/BMP2K probe known as SGC-AAK1-1 (Figure 1A).^22^ Since these compounds were known to be AAK1 active, the first step was to compare the cell-free binding affinity at 1 μM of each analog for AAK1 and CDKL2 (Figure 1B). The kinome-wide selectivity of each compound at 1 μM (S_10_(1 μM)) was considered in parallel (Figure 1C). The S_10_ score expresses selectivity and corresponds with the percent of the kinases screened that bind with a percent of control (PoC) value <10. When calculating a selectivity score, only the 403 wild-type human kinases in the DiscoverX *scanMAX* panel are considered. Binding affinity and kinome-wide selectivity scores for all analogs in Figure 1 are included in Table 1. Most of these compounds were exemplified in our campaign to identify an AAK1/BMP2K probe.^22^

**Table 1.** Potency and selectivity data for acylaminoindazole analogs.

| Compound | CDKL2 | AAK1 | $S_{10}(1 \mu\text{M})$ | # Kinases with | CDKL2 radiometric | CDKL2 NanoBRET |
| --- | --- | --- | --- | --- | --- | --- |

|  | PoC | PoC | score<sup>[a]</sup> | PoC <10<sup>[b]</sup> | enzymatic assay IC<sub>50</sub> (nM) | assay IC<sub>50</sub> (nM) |
| --- | --- | --- | --- | --- | --- | --- |
| SGC-AAK1-1 | 48 | 24 | 0.02 | 8 |  | >10000 |
| **1** | 65 | 18 | 0.017 | 7 |  |  |
| **2** | 11 | 30 | 0.002 | 1 |  |  |
| **3** | 28 | 20 | 0.01 | 4 |  |  |
| **4** | 22 | 38 | 0.022 | 9 |  |  |
| **5** | 41 | 47 | 0.005 | 2 |  |  |
| **6** | 25 | 33 | 0.007 | 3 |  | 1200 |
| **7** | 55 | 30 | 0.022 | 9 |  |  |
| **8** | 60 | 30 | 0.017 | 7 |  |  |
| **9** | 4.3 | 37 | 0.002 | 1 | 230 | 460 |
| **10** | 40 | 9.2 | 0.007 | 3 |  |  |
| **11** | 20 | 39 | 0.002 | 1 |  |  |
| **12** | 67 | 32 | 0.002 | 1 |  |  |
| **13** | 69 | 33 | 0 | 0 |  | >10000 |
| **14** | 97 | 47 | 0 | 0 |  | >10000 |
| **15** | 5 | 14 | 0.01 | 4 | 450 | 700 |
| **16** | NT<sup>[c]</sup> | NT | NT | NT |  | >10000 |

<sup>[a]</sup> S<sub>10</sub>(1 μM): percentage of screened kinases with PoC <10 at 1 μM; <sup>[b]</sup> PoC: percent of control values determined at 1 μM via DiscoverX *scan*MAX profiling; <sup>[c]</sup> NT: not tested.

**Figure 1.**
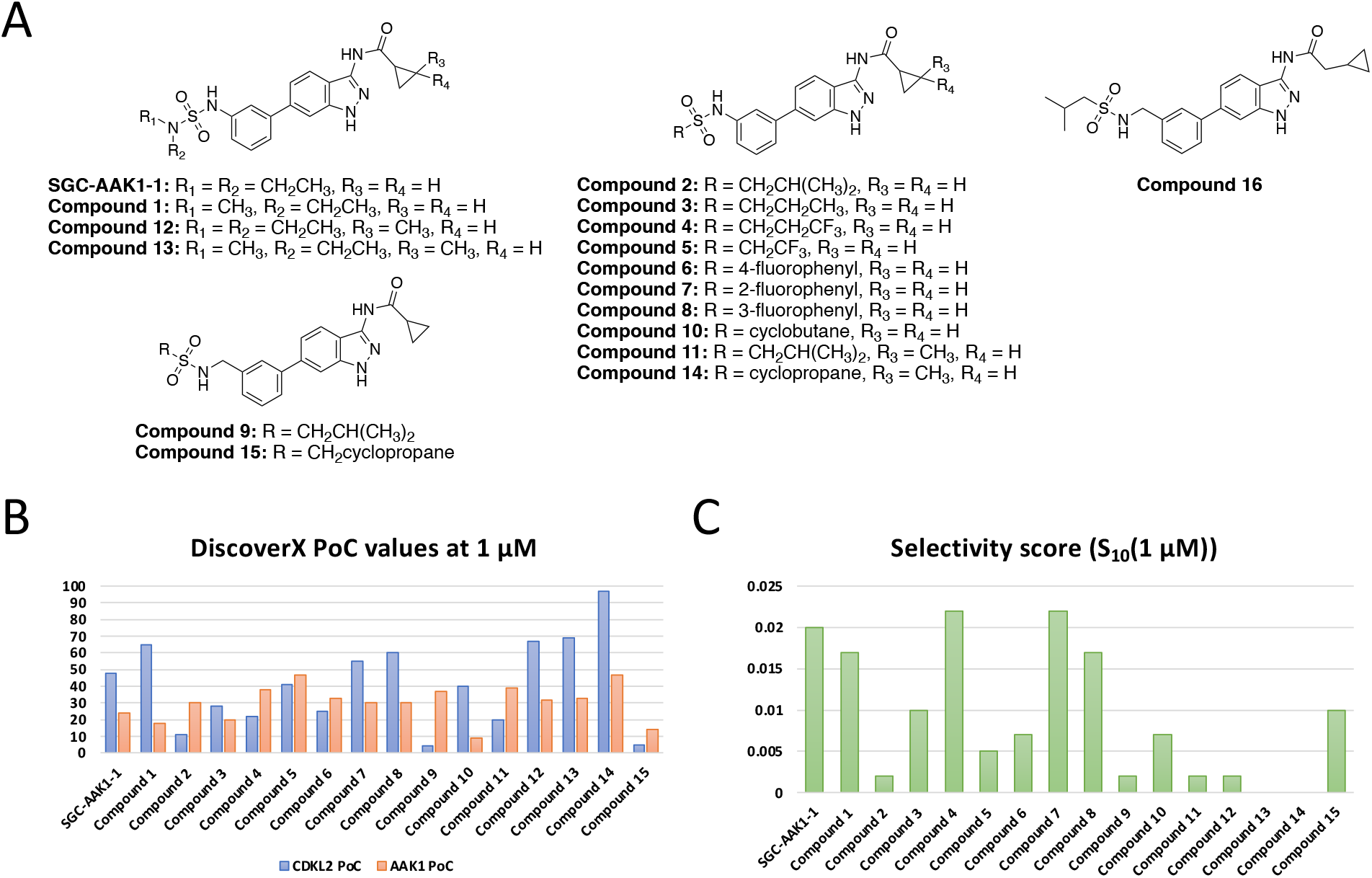
Structures and data for acylaminoindazole analogs. (A) Structures of all acylaminoindazole analogs considered herein. (B) Comparison of DiscoverX percent of control (PoC) values at 1 μM for CDKL2 and AAK1. (C) Selectivity score for each analog when analyzed versus 403 wild-type human kinases at 1 μM.

Examination of the data in Table 1 and Figure 1B demonstrated that nearly all compounds in the series demonstrated modest AAK1 binding affinity (PoC values 10–50 at 1 μM). Since this AAK1 affinity in the DiscoverX binding assay corresponded with sub-micromolar activity in the AAK1 NanoBRET assay,^22^ we hypothesized that most of these compounds are efficacious inhibitors of AAK1. Turning our attention to CDKL2 affinity, we noted that nine acylaminoindazole analogs bound to CDKL2 with PoC values >35. Of the remaining six analogs, only two demonstrated CDKL2 PoC values <10: compounds **9** and **15**. From a structural standpoint, these two analogs were unique from the other analogs in their inclusion of a methylene space between the phenyl ring and sulfonamide (Figure 1A). While a larger window was observed between the AAK1 and CDKL2 affinity for compound **9** and the kinome-wide selectivity of compound **9** was better than that of **15**, both analogs were prioritized for follow-up.

As shown in Table 1 and Figure 2, both compounds inhibited CDKL2 in a radiometric enzymatic assay with IC_50_ values <500 nM. Given this confirmed functional inhibition, compounds **9** and **15** were next evaluated in the CDKL2 NanoBRET assay to gauge their ability to engage CDKL2 in cells. Three replicates of the CDKL2 NanoBRET assay for compound **9** were averaged to an IC_50_ value of 460 nM (Figures 2 and S1). The IC_50_ value for compound **15** in the CDKL2 NanoBRET assay was found to be 700 nM (Table 1 and Figure S2). The in-cell potency of compound **9** motivated a deeper dive to confirm its kinome-wide selectivity. As shown in Figure 2B, when broadly profiled against 403 wild-type human kinases compound **9** only bound one kinase with PoC <10, supporting its selectivity score (S_10_(1 μM) = 0.002) in Table 1. Nine additional kinases demonstrated a PoC <40 in this broad profiling effort (Figure 2C). To determine whether binding resulted in inhibition of kinase activity, radiometric enzymatic assays executed at the Km value for ATP for each respective kinase were used to evaluate compound **9** versus all kinases that bound with PoC <40 at 1 μM (Figure 2C). This effort narrowed the kinases inhibited by compound **9** to only three: CDKL2, BMP2K, and AAK1. We previously documented the activity of acylaminoindazoles on AAK1 and BMP2K^22^ and this activity was maintained for compound **9** in cell-free assays. Next, the cellular target engagement of AAK1 and BMP2K by compound **9** was evaluated. This compound was found to have higher affinity for CDKL2 than it does for AAK1 and BMP2K in cells (Figure 2C). Truncated versions of AAK1 (1–353) and BMP2K (1–367) and full-length CDKL2 were employed in the respective radiometric enzymatic assays (Figure 2C). Full-length versions of all three kinases were used in the NanoBRET assays (Figure 2C). The difference in the length of protein used for AAK1 and BMP2K in each assay could, in part, explain the distinct results observed in the radiometric enzymatic versus NanoBRET assays. In contrast, full-length CDKL2 protein was employed in both radiometric enzymatic and NanoBRET assays and results when compound **9** was evaluated were more consistent between them.

**Figure 2.**
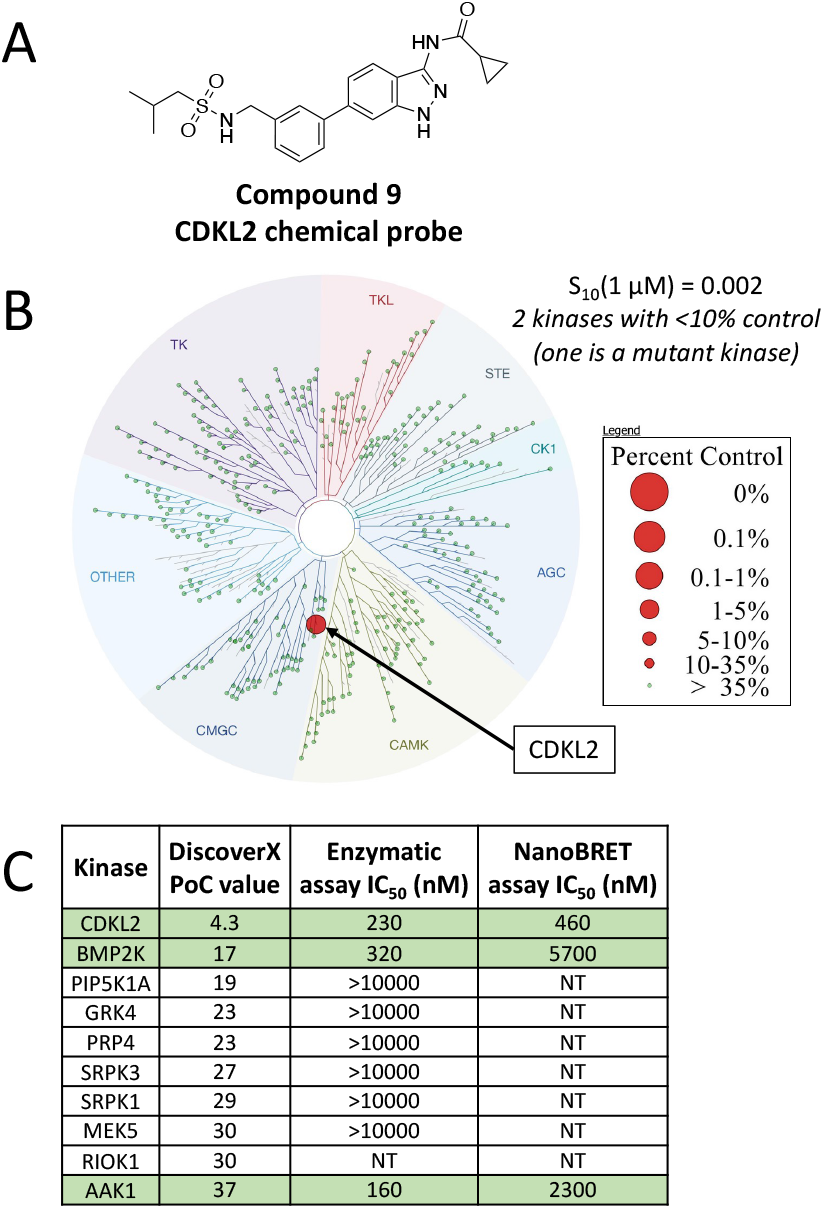
Structure, potency, and kinome-wide selectivity data related to CDKL2 chemical probe (compound **9**). (A) Structure of compound **9**. (B) Kinome dendrogram that illustrates non-mutant kinases that bind with PoC <10 when compound **9** was screened at 1 μM in the DiscoverX *scanMAX* panel. (C) Data for all human wild-type kinases in the DiscoverX *scanMAX* panel that bound with a PoC <40 when compound **9** was screened at 1 μM (column 2), enzymatic IC_50_ values generated when compound **9** was evaluated in each kinase radiometric enzymatic assay (column 3), and NanoBRET IC_50_ values generated when compound **9** was evaluated in each kinase NanoBRET assay (column 4). PoC = percent of control.

A deeper look was next taken at the selectivity of compound **9** within the CDKL family. The PoC data for most of this family was collected when compound **9** was profiled in the DiscoverX *scanMAX* panel at 1 μM (Figure 3A). Compound **9** only demonstrated appreciable binding affinity to CDKL2 in this set of assays. The kinase domains of CDKL1, CDKL2, and CDKL3 were next employed in thermal shift assays with increasing concentrations of compound **9**. A dose-dependent change in the melting temperature (ΔTm) of CDKL2 was observed in the presence of this compound, while the ΔTm of neither CDKL1 nor CDKL3 was perturbed by compound **9** (Figure 3B). Finally, *in vitro* kinase assays, using the ADP-Glo assay to readout inhibition of activity, were executed to evaluate whether compound **9** inhibits the activity of human and/or mouse forms of CDKL family members (Figure 3C). Substrate phosphorylation was not inhibited at concentrations up to 10 μM of compound **9** for CDKL1, CDKL3, CDKL4, or CDKL5. Inhibition of the kinase activity of CDKL2 by compound **9**, however, was observed. Substrate phosphorylation by the human and mouse forms of CDKL2 was inhibited with IC_50_ values of 43 nM and 21 nM, respectively. The perceived shift in IC_50_ values between the CDKL2 radiometric enzymatic assay (Figure 2C) and the CDKL2 *in vitro* kinase assays (Figure 3C) could be due, in part, to the difference in concentration of ATP used in each assay, 200 μM for the radiometric enzymatic assay and 50 μM for the *in vitro* ADP-Glo kinase assay. Slightly different constructs were also used in each assay. While both assays measure the ability of a compound to inhibit phosphorylation of a substrate by full-length CDKL2, the radiometric enzyme assay specifically probes transfer of a radiolabeled gamma phosphate from ATP to a peptide substrate and the ADP-Glo assay assesses the conversion of ATP to ADP as a measure of kinase activity. The higher concentration of ATP used in the radiometric enzymatic assay provides more competition for binding to the same site as compound **9**, resulting in a comparatively elevated IC_50_ value in this assay. Overall, however, the trends are the same and compound **9** is a verified, potent inhibitor of CDKL2 activity.

**Figure 3.**
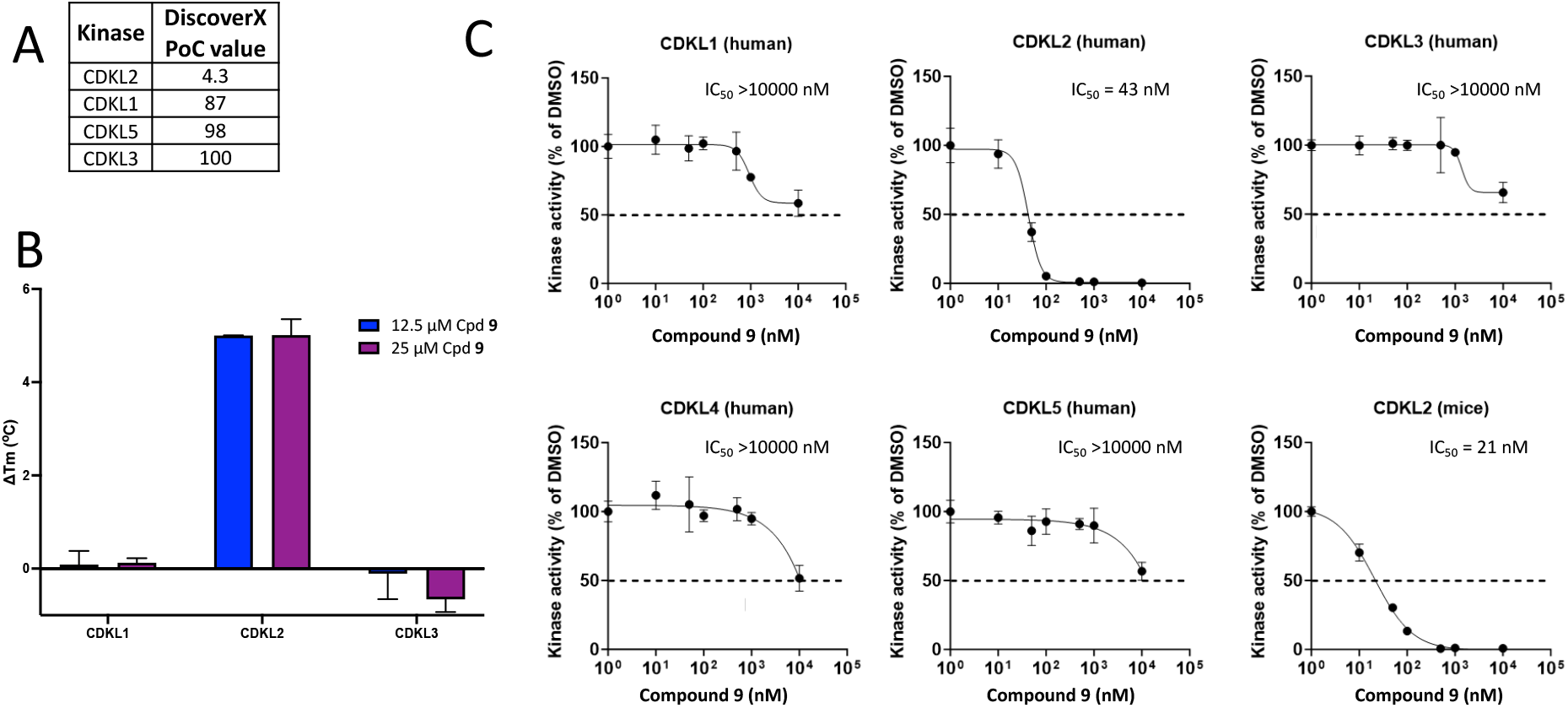
Data that supports the selectivity of CDKL2 chemical probe (compound **9**) within the CDKL family. (A) DiscoverX PoC values generated using the available CDKL family binding assays and screening at 1 μM of compound **9**. (B) Thermal shift assay data for compound **9** when evaluated versus CDKL1, CDKL2, and CDKL3 kinase domains. (C) *In vitro* kinase assay results for compound **9** when evaluated versus full-length human CDKL1–5 and murine CDKL2 and the ADP-Glo assay was used as a readout. PoC = percent of control; Cpd = compound.

With a CDKL2 probe candidate identified within the acylaminoindazole series we next sought a suitable negative control compound. To meet criteria, a negative control candidate must be structurally similar to the chemical probe but lack CDKL2 inhibitory activity. A combination of the CDKL2 affinity in the DiscoverX assay and kinome-wide selectivity was considered when choosing potential candidates (Table 1). Accordingly, compounds **6, 13, 14**, and SGC-AAK1-1 were analyzed in the CDKL2 NanoBRET assay. Compound **6** was found to have micromolar affinity for CDKL2 in cells and was excluded from consideration (Figure S2). Gratifyingly, SGC-AAK1-1 was found to demonstrate an IC_50_ >10 μM for CDKL2 in cells (Figure S2). This makes SGC-AAK1-1 a complementary molecule to use in concert with compound **9** to tease apart cellular phenotypes resulting from AAK1/BMP2K inhibition versus those from CDKL2 inhibition when compound **9** is dosed at concentrations exceeding 1 μM. While compounds **13** and **14** were found to lack CDKL2 affinity in cells (Figure S2), the chiral center on the cyclopropane ring made us less enthusiastic about these compounds and motivated preparation of a more suitable negative control based upon our established structure–activity relationships. Compound **16** (Figure 1) was next designed as a putative negative control compound. When compared to compound **9**, only one part of the molecule was modified. A methylene spacer was placed between the carbonyl and cyclopropyl ring on compound **9** to furnish compound **16**. Once prepared, compound **16** was evaluated in the CDKL2, AAK1, and BMP2K NanoBRET assays. It was found to lack affinity for all these kinases (IC_50_ >10 μM, Figure S3) and thus was selected as an appropriate negative control, differing only by a single methylene, to be used alongside compound **9**.

Structural studies were employed to understand the binding mode and rationalize the selectivity of compound **9** for CDKL2. A co-crystal structure of compound **9** bound to CDKL2 was solved (Figure 4, Table S1) and interactions mapped using Protein–Ligand Interaction Profiler (PLIP).^23^ This structure shows a collapsed P-loop enveloping and stabilizing the binding of compound **9** and an αC-out, DFG-in type conformation. While the compounds are structurally very different, this binding mode is similar to the previously discussed crystal structure with TSC 2312 (PDB code: 4BBM) and distinct from the CDKL2 structure with CDK1/2 Inhibitor III (PDB code: 4AAA).^21^ Like TSC 2312, compound **9** harnesses the inactive αC-out conformation that is unique to CDKL2 versus other CDKL family members.

**Figure 4.**
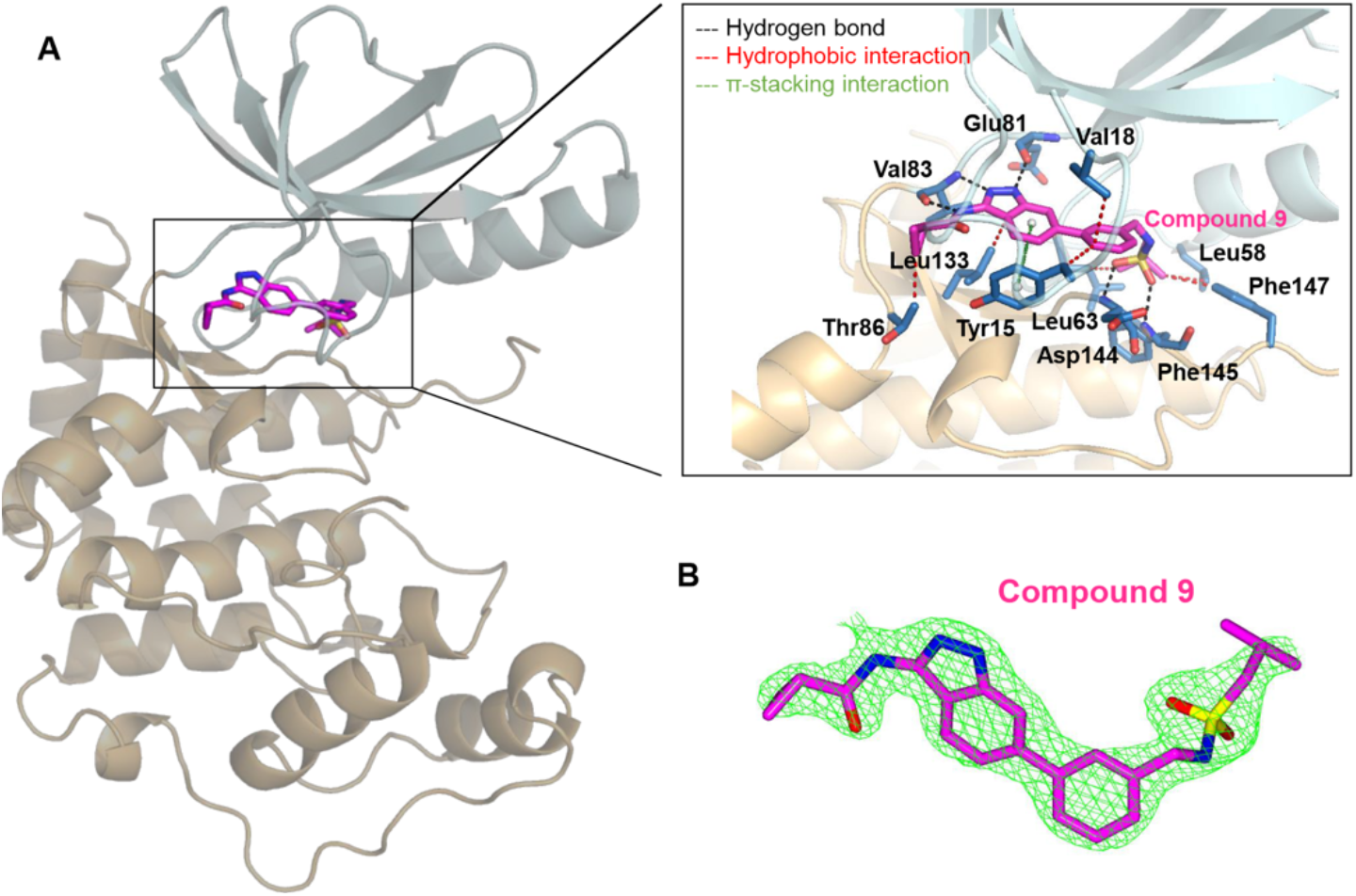
Co-crystal structure of CDKL2 with compound **9** (PDB code: 8S6I) (A) Overview of CDKL2 with compound **9**. N- and C-terminal lobes colored in pale cyan and light orange, respectively. Compound **9** (magentas sticks) binds to the active site between the two lobes. (Inset) Interactions of compound **9** with CDKL2. Black dashed lines: hydrogen bonds (Glu81, Val83, Asp144, and Phe145); red dashed lines: hydrophobic interactions (Tyr15, Val18, Leu58, Leu63, Thr86, Leu133, and Phe147); green dashed lines: π-stacking interaction (Tyr15). Interactions mapped using Protein–Ligand Interaction Profiler (PLIP). (B) mFo-DFc omit electron density map for compound **9**. The green mesh represents electron density of compound **9** at 2.0 σ (CCP4 molecular graphics).

Comparison of the structure of the AAK1/BMP2K probe bound to BMP2K (PDB code: 5I3R) to this new CDKL2 co-crystal structure shows that the conserved nitrogens of the acylaminoindazole core make key hydrogen bonds with Glu131 and Cys133 in BMP2K^22^ and Glu81 and Val83 in CDKL2. These hydrogen bonds anchor the two compounds in similar orientations to the kinase hinge region in the respective active sites. The pendant aryl ring of compound **9** is accommodated by a large hydrophobic region of the ATP-binding site, which is common to BMP2K as well.^22^ Compound **9** bears a sulfonamide that hydrogen bonds with Phe145 and Asp144, which are part of the DFG motif. The methylene that has been inserted between the pendant aryl ring and sulfonamide is essential in the placement of this sulfonamide in proximity to the key residues mentioned and for hydrophobic interactions with Phe147. This importance of the methylene is supported by the CDKL2 affinity of compounds **9** and **15** versus that of analogs that lack the methylene spacer in our aminoindazole library (Table 1). Finally, comparison of the BMP2K and CDKL2 structures helps explain the selectivity of compound **9** for CDKL2 versus AAK1 and BMP2K. The larger hydrophobic pocket in CDKL2 can enclose the longer sulfonamide side chain whereas the same pocket in BMP2K, and AAK1 by analogy given the highly homologous sequence, is smaller and would introduce steric clash when compound **9** is bound. The ATP-binding sites of other kinases are similarly unlikely to accommodate this sulfonamide, precluding their binding to compound **9** with high affinity and imparting the kinome-wide selectivity we observe.

The hydrophobic pocket that accommodates the sulfonamide, however, is large. Thus, analogs with smaller alkyl sulfonamides (**2**–**5** and **10**) bind in the DiscoverX CDKL2 assay (Table 1, CDKL2 PoC values 11–41). These compounds are proposed to not encounter steric clash but also not benefit from the key hydrogen bonds with Glu81 and Val83 because they are positioned too far from these residues. Compounds bearing an aryl sulfonamide with an *ortho*-(**7**) or *meta*-fluorine (**8**) are weaker binders (Table 1, CDKL2 PoC 55–60), while the *para*-fluorinated phenyl sulfonamide (**6**) demonstrates slightly higher affinity for CDKL2 (Table 1, CDKL2 PoC 25). The pocket tolerates these larger rings and the DiscoverX CDKL2 PoC values support that the *para*-fluorine may be positioned to make additional interactions that the *ortho*- and *meta*-fluorine cannot. Finally, SGC-AAK1-1 and compound **1** are weaker binders to CDKL2 (Table 1, CDKL2 PoC 48–65). Since the sulfonamides on these two compounds are not significantly larger than those on other analogs, the added nitrogen and/or the branching (two R groups) could lead to unfavorable interactions with the binding pocket. The solved structure of compound **9** shows that the space around the cyclopropane ring in compound **9** is limited. Thus, modification of and near the cyclopropane ring is likely to perturb key interactions with Glu81 and Val83 due to steric clash and forced repositioning of analogs in the CDKL2 ATP-binding site. This finding provides rationale for why analogs bearing a methylated cyclopropyl ring (**11, 12, 13, 14**) and the deeper projecting negative control (**16**) lack affinity for CDKL2.

With a potent and selective CDKL2 chemical probe in hand, we next evaluated its impact on downstream signaling. The Ultanir lab recently reported that CDKL2 phosphorylates microtubule end binding protein 2 (EB2) at Ser222 in HEK293T cells, primary neurons, and *in vivo*.^24^ This finding motivated examination of the response of rat primary neurons to treatment with compound **9**. It is important to note, however, that CDKL5 is responsible for approximately 80% of EB2 phosphorylation.^24^ Because CDKL5 is not inhibited by our compound, we expected to see a maximal 20–25% reduction of EB2 phosphorylation by the CDKL2 probe. We evaluated a CDKL5 chemical probe (SGC-CAF382-1) in parallel at a concentration (50 nM) where ∼80% reduction in EB2 phosphorylation was previously observed.^25^ Neurons were treated with increasing concentrations of compound **9** alone or 50 nM of SGC-CAF382-1 with increasing concentrations of compound **9** to explore whether these two compounds could completely inhibit EB2 phosphorylation when dosed together. Negative control compound **16** was evaluated in parallel to explore chemotype-induced effects on EB2 phosphorylation. As shown in Figures 5 and S4, compound **9** completely suppressed the CDKL2-mediated EB2 phosphorylation in a dose-dependent manner following 1h exposure of rat primary neurons. Total EB2 expression was not impacted. In contrast, negative control compound **16** did not perturb EB2 phosphorylation or total EB2 expression at concentrations up to 10 μM. The concentration at which EB2 phosphorylation is inhibited by compound **9** corresponds well with its IC_50_ value in the CDKL2 NanoBRET assay (460 nM). This result confirmed that CDKL2 inhibition by compound **9** disrupts CDKL2-mediated downstream signaling in rat primary neurons. Treatment of rat neurons with SGC-CAF382-1 alone resulted in a robust reduction in EB2 phosphorylation (∼95%). The co-dosing experiments with SGC-CAF382-1 confirmed that EB2 phosphorylation can be completely suppressed (∼98.5%) when CDKL5 and CDKL2 are both inhibited.

**Figure 5.**
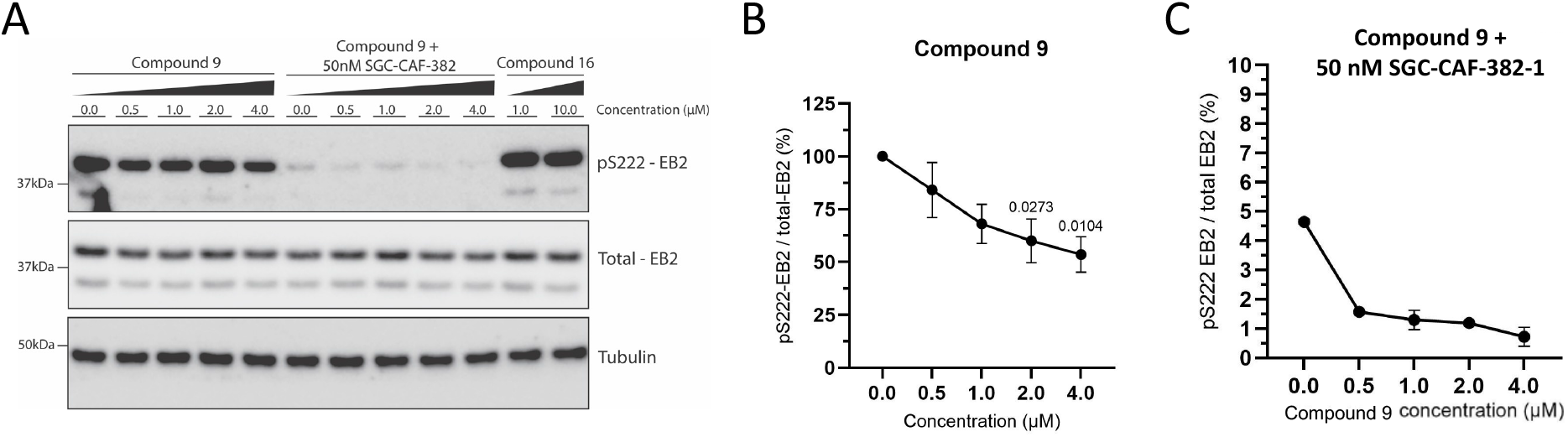
Western blot analyses of pS222-EB2 in rat primary neurons after 1 h treatment with compound **9**, SGC-CAF382-1, compound **9** + SGC-CAF382-1, or compound **16**. (A) Representative blot from a single replicate, replicate two in Figure S4. (B) Quantification of pS222-EB2 normalized to total EB2 for compound **9** treatment, n = 4. Error bars represent SEM. One way ANOVA test: p-value = 0.0191. (C) Quantification of pS222-EB2 normalized to total EB2 for compound **9** + SGC-CAF382-1 treatment, n = 2. Error bars represent SEM.

As mentioned, CDKL2 is an identified regulator of epithelial–mesenchymal transition (EMT), a process that can accelerate cancer progression.^5^ It was reported that shRNA directed at CDKL2 in breast cancer, specifically human mammary gland epithelial (HMGE) cells, decreased protein expression of vimentin and N-cadherin and mRNA levels of ZEB1 and CD44.^5^ Rather than HMGE cells, we selected an epithelial (MCF-7) and mesenchymal (MDA-MBA-231) breast cancer cell line with known CDKL2 expression at the mRNA level^5^ to probe whether these same responses could be recapitulated using our CDKL2 chemical probe. When these cells were treated in dose–response format with 1, 5, or 10 μM of compound **9** for 24, 48, or 72 hours, no reproducible impact was observed with respect to the protein expression of CD44, ZEB1, or β-catenin (Figure S5). Furthermore, when exposed to the same concentrations and time course, protein expression of E-cadherin was not altered in MCF-7 cells and vimentin in MDA-MB-231 cells. Results were inconsistent and often compounds **9** and **16** elicited a nearly identical response in these studies. We were not able to pharmacologically modulate expression of the selected EMT proteins using compound **9** at concentrations where it is CDKL2 active in cells based on NanoBRET and western blot data. At this biologically relevant concentration, we also noted that CDKL2 inhibition by compound **9** did not impact the viability of MCF7 or MDA-MB-231 cells when treated for 48 hours (Figure S6). Furthermore, a similar lack of toxicity at relevant concentrations was observed when a patient derived xenograft (PDX) model of triple-negative breast cancer (TU-BcX-4IC cells)^26^ was treated in dose–response format with compound **9** (Figure S6). TU-BcX-4IC cells were originally derived from a patient with metaplastic triple-negative breast carcinoma, making it one of the most aggressive and dangerous subtypes of breast cancer. Versus immortalized cell lines, PDX-derived cell lines more accurately mimic the behavior of actual tumor cells in patients.^27-29^ These TU-BcX-4IC cells did not display significant changes in morphology, total area, or average cell size at 1 μM (Figure S6). MDA-MB-231 cells also did not demonstrate changes in these features when treated at 1 μM (data not shown).

We have described the evaluation of a series of acylaminoindazoles as inhibitors of CDKL2. Analyses of a plethora of binding and kinome-wide selectivity data for this set of compounds enabled selection of compound **9** as our CDKL2 chemical probe. This compound demonstrated inhibition of CDKL2 enzymatic activity in cell-free assays and nearly equivalent engagement of CDKL2 in cells. Extensive selectivity screening, including evaluation of binding and inhibition of putative off-target kinases, has verified that compound **9** inhibits very few kinases. In enzymatic assays, this compound inhibits CDKL2, AAK1, and BMP2K with nearly equivalent IC_50_ values. In cells, however, it is a higher affinity binder to CDKL2 and demonstrates a modest 5–12-fold enhanced binding affinity for CDKL2 when compared to AAK1 and BMP2K. This provides a cautionary note as well, that compound **9** should be used at concentrations of ≤1 μM in cells to avoid perturbation of AAK1 and BMP2K. It is also strongly suggested that our designed and confirmed negative control, compound **16**, and SGC-AAK1-1 are used in tandem with compound **9** in experiments designed to probe CDKL2-mediated biology. Structural studies helped illuminate key interactions of our CDKL2 chemical probe with the ATP-binding site and rationalize potency trends within the acylaminoindazole series. Comparison of the CDKL2 co-crystal structure with compound **9** to that of SGC-AAK1-1 bound to BMP2K generated hypotheses about the selectivity of compound **9** for CDKL2 when screened broadly. Inhibition of downstream signaling without an associated impact on viability was observed when rat primary neurons were treated with compound **9**, supporting that binding to CDKL2 in cells impacted activity of the kinase. A similar lack of changes in viability were noted when breast cancer cells were treated with the probe pair. Moreover, we found that compound **9** does not phenocopy CDKL2 shRNA results related to EMT and does not alter the expression of the specific proteins involved in this process in the cell lines, concentrations, and time points that we probed.

## ASSOCIATED CONTENT

### Supporting Information

Supplemental figures (synthetic schemes, NanoBRET assay curves, western blot replicates, viability results), a table (X-ray crystallographic data), and experimental details, including synthetic schemes and characterization of key compounds, protocols related to selectivity screening and biological assays, and crystallographic details.

### Accession Codes

The PDB accession code for the X-ray co-crystal structure of CDKL2 + **9** is 8S6I.

## AUTHOR INFORMATION

### Notes

The authors declare no competing financial interest.

## ACKNOWLEDGMENT

Promega kindly provided constructs for NanoBRET measurements of CDKL2, AAK1, and BMP2K. The kinome tree in Figure 2 was prepared using the TREE*spot* kinase interaction mapping software at http://treespot.discoverx.com. We acknowledge the Department of Chemistry Mass Spectrometry Core Laboratory at the University of North Carolina for assisting with mass spectrometry analyses. We thank the beamline scientists at the Swiss Light Source (PSI) for their great support during data collection.

The Structural Genomics Consortium (SGC) is a registered charity (number 1097737) that receives funds from Bayer AG, Boehringer Ingelheim, the Canada Foundation for Innovation, Eshelman Institute for Innovation, Genentech, Genome Canada through Ontario Genomics Institute, EU/EFPIA/OICR/McGill/KTH/Diamond, Innovative Medicines Initiative 2 Joint Undertaking (EUbOPEN grant number 875510), Janssen, Merck KGaA (aka EMD in Canada and USA), Pfizer, the São Paulo Research Foundation-FAPESP, and Takeda. Research reported in this publication was supported in part by NC Biotechnology Center Institutional Support Grant 2018-IDG-1030 and NIH U24DK116204. Ultanir lab was supported by the Francis Crick Institute which receives its core funding from Cancer Research UK (CC2037), the UK Medical Research Council (CC2037), and the Wellcome Trust (CC2037).

## Abbreviations

IC_50_: half maximal inhibitory concentration
NanoBRET: bioluminescence resonance energy transfer using NanoLuciferase
nLuc: NanoLuciferase
nM: nanomolar
PoC: percent of control
PDB: protein data bank
SEM: standard error of the mean.

