## Supplemental Information for "Discovery and Characterization of a Chemical Probe for Cyclin-Dependent Kinase-Like 2"

### Table of Contents


| | |
|---|---|
| Chemistry | S2 |
| Scheme 1 | S3 |
| Scheme 2 | S3 |
| Scheme 3 | S4 |
| Scheme 4 | S4 |
| *In Vitro* Studies | S5 |
| Figure S1 | S8 |
| Figure S2 | S9 |
| Figure S3 | S9 |
| Figure S4 | S9 |
| Figure S5 | S10 |
| Figure S6 | S11 |
| Crystallography Methods | S12 |
| Table S1 | S12 |
| Purity traces and spectra for all compounds | S14–S21 |
| References | S22 |

#### CHEMISTRY

##### General Information.

No unexpected or notable safety hazards occurred in performing the chemistry described below. Reagents and solvents were used as received from commercial suppliers without purification. Reactions were executed under argon atmosphere and at ambient temperature (~25 °C) unless otherwise indicated. A rotary evaporator was used to remove solvent under reduced pressure. Thin layer chromatography (TLC) and LC–MS were employed to track reaction progress both before and after purification. The following abbreviations are used in the schemes and/or experimental procedures that follow: °C (degrees Celsius), mmol (millimoles), μmol (micromoles), mL (milliliters), μL (microliters), mg (milligrams), eq (equivalent(s)), h (hours), min (minutes), M (molar), N (normal), HRMS (high resolution mass spectrometry), ESI (electrospray ionization), *m*/*z* (mass/charge number), LC–MS (liquid chromatography–mass spectrometry), HPLC (high-performance liquid chromatography), C (carbon), and H (hydrogen). Common reagent abbreviations include: DIPEA (*N, N*-diisopropylethylamine), DMF (*N, N*-dimethylformamide), DMSO (dimethyl sulfoxide), EDC (1-ethyl-3-(3-dimethylaminopropyl)carbodiimide), EtOAc (ethyl acetate), HCl (hydrochloric acid), HOBt (N-hydroxybenzotriazole), MeOH (methanol), and TFA (trifluoroacetic acid). Reported yields correspond with isolated, pure product.

Nuclear magnetic resonance (NMR) spectra and microanalytical data were gathered for intermediates and final compounds to corroborate their identity and evaluate their purity. $^{1}H$ and $^{13}C$ NMR spectra were recorded in DMSO-$d_6$ or methanol-$d_4$ on a Bruker AVANCE III 850 megahertz (MHz), Bruker AVANCE NEO 500 MHz, or Bruker AVANCE NEO NANO 400 MHz spectrometer. Chemical shifts are line listed in parts per million (ppm) with deuterated solvent peaks labeled and referenced as the internal standard. Coupling constants ($J$ values) have been calculated and are listed in hertz (Hz). Spin multiplicities for peaks include: singlet (s), doublet (d), doublet of doublets (dd), doublet of pentets (dp), triplet (t), doublet of triplets (dt), and multiplet (m). MestReNova was used to process all NMR data.

High-resolution mass spectrometry (HRMS) samples were analyzed via a ThermoFisher Q Exactive HF-X mass spectrometer coupled with a Waters Acquity H-class liquid chromatograph system. Heated electrospray source (HESI) was used to deliver 3 μL of sample compounds at a flow rate of 0.3 mL/minute. These solutions were prepared at a maximal concentration of 0.1 mg/mL The electrospray source conditions were set as follows: spray voltage 3.0 kV, sheath gas (nitrogen) 60 arb, auxiliary gas (nitrogen) 20 arb, sweep gas (nitrogen) 0 arb, nebulizer temperature 375°C, capillary temperature 380°C, and RF funnel 45 V. A mass range of 150–2000 m/z was sampled and measurements were recorded at a resolution of 120,000. A Waters Acquity UPLC BEH C18 column (2.1 x 50 mm, 1.7 um particle size) was employed for separations, using 95% water with 0.1% formic acid ramped linearly over 5 minutes to 100% acetonitrile with 0.1% formic acid and held until 6 minutes. At 7 minutes, the gradient was switched back to 95% water with 0.1% formic acid and allowed to re-equilibrate until 9 minutes. Xcalibur (ThermoFisher) was used to analyze all data. Molecular Formula Calculator (v 1.2.3) was used to determine molecular formulas. All observed species were singly charged, as verified by unit *m/z* separation between mass spectral peaks corresponding to the $^{12}C$ and $^{13}C^{12}C_{c-1}$ ion contributions for each elemental composition. Purity was determined using LC–MS and/or NMR spectroscopy. All intermediates and final compounds demonstrated >95% purity by these analyses.

#### Synthetic methods for key compounds.

**Scheme S1.** Synthetic route for preparation of intermediate **17**.

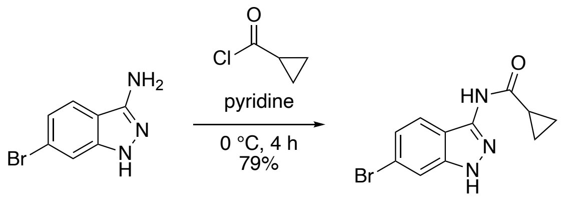

*N-(6-bromo-1H-indazol-3-yl)cyclopropanecarboxamide* (**17**). To a suspension of 6-bromo-1*H*-indazol-3-amine (500 mg, 1 eq, 2.36 mmol) in pyridine (7.0 mL) was added cyclopropanecarbonyl chloride (246 mg, 214 µL, 1 eq, 2.36 mmol) dropwise at 0 °C. The resulting mixture was stirred at 0 °C for 4 h. The reaction solution was added 4 mL of deionized water, and the resulting mixture was concentrated *in vacuo*. The residue was dissolved in 4 mL of DMF, and deionized water (15 mL) was added. The precipitate that formed was filtered, washed with water, and dried under vacuum to afford compound **17** as a colorless solid (525 mg, 79% yield). $^{1}$H NMR (400 MHz, DMSO-$d_6$) δ 12.73 (s, 1H), 10.74 (s, 1H), 7.76 (d, $J$ = 8.7 Hz, 1H), 7.64 (d, $J$ = 1.7 Hz, 1H), 7.15 (dd, $J$ = 8.7, 1.7 Hz, 1H), 1.96 – 1.88 (m, 1H), 0.88 – 0.79 (m, 4H). $^{13}$C NMR (214 MHz, DMSO-$d_6$) δ 174.97, 144.84, 143.96, 127.73, 125.56, 123.03, 118.09, 115.65, 16.77, 10.48.

**Scheme S2.** Synthetic route for preparation of compound **9**.

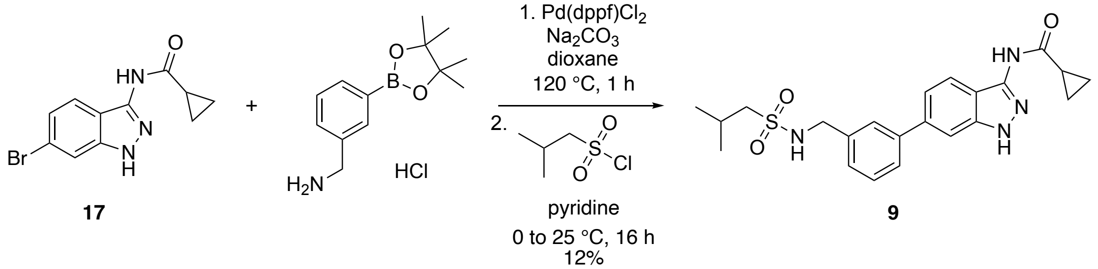

*N-(6-(3-(((2-methylpropyl)sulfonamido)methyl)phenyl)-1H-indazol-3-yl)cyclopropanecarboxamide* (**9**). To a microwave vial was added *N*-(6-bromo-1*H*-indazol-3-yl)cyclopropanecarboxamide (50 mg, 1 eq, 0.18 mmol), (3-(4,4,5,5-tetramethyl-1,3,2-dioxaborolan-2-yl)phenyl)methanamine hydrochloride (48 mg, 1 eq, 0.18 mmol), and Pd(dppf)$Cl_2$ (13 mg, 0.1 eq, 18 µmol) in a mixture of 1,4-dioxane (0.8 mL) and 1M aqueous sodium carbonate (0.4 mL). The vial was capped and heated in the microwave at 120 °C for 30 min. Once cool, the reaction progress was examined by TLC, which revealed a fair amount of residual starting material. The reaction mixture was treated with Pd(dppf)$Cl_2$ (13 mg, 0.1 eq, 18 µmol), capped, and heated in the microwave at 120 °C for another 30 min. The mixture was filtered through a thin pad of Celite and washed with EtOAc. The filtrate was washed with brine, dried over $Na_2SO_4$, filtered, and concentrated. The crude reaction mixture was taken to the next step with no further purification.

To a cooled solution of crude *N*-(6-(3-(aminomethyl)phenyl)-1*H*-indazol-3-yl)cyclopropanecarboxamide (55 mg, 1 eq, 0.18 mmol) in pyridine (1.5 mL) was added dropwise isobutylsulfonyl chloride (56 mg, 47 µL, 2 eq, 0.36 mmol) at 0 °C. The reaction mixture was stirred at room temperature overnight. Solvent was removed and the residue was partitioned between EtOAc and saturated aqueous sodium bicarbonate solution. The organic layer was dried over $Na_2SO_4$, filtered, and concentrated. The residue was purified by flash column chromatography

($SiO_2$, 0−5% MeOH in $CH_2Cl_2$) to yield **9** (9.2 mg, 12% yield) as a colorless solid. $^1$H NMR 500 MHz, DMSO-*d*$_6$) δ 12.69 (s, 1H), 10.68 (s, 1H), 7.87 (d, *J* = 8.5 Hz, 1H), 7.71 (t, *J* = 2.0 Hz, 1H), 7.67 (t, *J* = 6.3 Hz, 1H), 7.63 (dt, *J* = 8.0, 1.5 Hz, 1H), 7.61 (s, 1H), 7.46 (t, *J* = 7.5 Hz, 1H), 7.38 – 7.31 (m, 2H), 4.24 (d, *J* = 5.5 Hz, 2H), 2.84 (d, *J* = 6.5 Hz, 2H), 2.06 (dp, *J* = 13.0, 6.5 Hz, 1H), 1.97 – 1.92 (m, 1H), 0.96 (d, *J* = 7.0 Hz, 6H), 0.88 – 0.83 (m, 4H). $^{13}$C NMR (214 MHz, DMSO-*d*$_6$) δ 174.86, 144.86, 143.71, 143.65, 142.34, 141.61, 132.15, 130.03, 129.65, 129.12, 126.35, 122.10, 118.60, 110.59, 62.42, 49.04, 27.37, 25.36, 16.80, 10.38, 3.25. HRMS (ESI): *m/z* calculated for $C_{22}H_{27}N_4O_3S$ [M+H]$^+$: 427.17256. Found: 427.18039.

**Scheme S3.** Synthetic route for preparation of intermediate **18**.

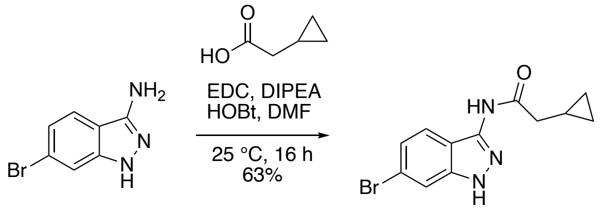


**18**

*N-(6-bromo-1H-indazol-3-yl)-2-cyclopropylacetamide* (**18**). To a solution of 2-cyclopropylacetic acid (56.7 mg, 55.1 µL, 1.2 eq, 566 µmol) in DMF (3 mL) was added 6-bromo-1*H*-indazol-3-amine (100 mg, 1 eq, 472 µmol), DIPEA (183 mg, 246 µL, 3 eq, 1.41 mmol), EDC (110 mg, 1.5 eq, 707 µmol), and HOBt (108 mg, 1.5 eq, 707 µmol). The reaction mixture was stirred at room temperature for 16 h. Solvent was removed, the residue dissolved in EtOAc and then washed with water, 2 N HCl, and brine. The organic layer was dried over $Na_2SO_4$, filtered, and concentrated. The residue was purified by flash column chromatography ($SiO_2$, 0−20% EtOAc in hexanes) to yield **18** (88 mg, 63% yield) as a colorless solid. $^1$H NMR (400 MHz, methanol-*d*$_4$) δ 8.48 (dd, *J* = 1.6, 0.4 Hz, 1H), 7.67 (dd, *J* = 8.4, 0.4 Hz, 1H), 7.45 (dd, *J* = 8.4, 1.6 Hz, 1H), 2.88 (d, *J* = 7.2 Hz, 2H), 1.26 – 1.15 (m, 1H), 0.58 – 0.54 (m, 2H), 0.26 (dt, *J* = 6.4, 4.4 Hz, 2H). $^{13}$C NMR (101 MHz, methanol-*d*$_4$) δ 173.94, 154.16, 141.85, 128.08, 124.98, 122.67, 120.50, 119.52, 40.88, 7.85, 4.84. Purity (LC–MS): 100%.

**Scheme S4.** Synthetic route for preparation of compound **16**.

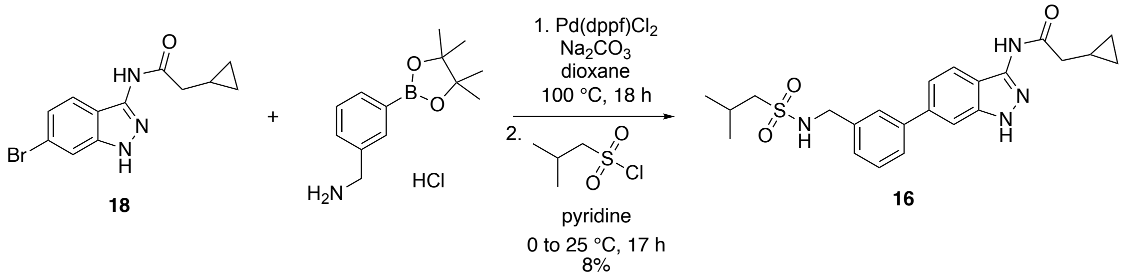


*2-cyclopropyl-N-(6-(3-(((2-methylpropyl)sulfonamido)methyl)phenyl)-1H-indazol-3-yl)acetamide* (**16**). To a microwave vial was added *N*-(6-bromo-1H-indazol-3-yl)-2-cyclopropylacetamide (50 mg, 1 eq, 0.17 mmol), (3-(aminomethyl)phenyl)boronic acid hydrochloride (32 mg, 1 eq, 0.17 mmol), and Pd(dppf)$Cl_2$ (25 mg, 0.2 eq, 34 µmol) in a mixture of 1,4-dioxane (1.0 mL) and 1 M sodium carbonate (0.5 mL). The reaction mixture was degassed and heated at 100 °C for 18 h. The mixture was filtered through a thin pad of Celite and washed with EtOAc. The filtrate was washed with brine, dried over $Na_2SO_4$, filtered, and concentrated. The crude reaction mixture was taken to the next step with no further purification.

To a cooled solution of crude *N*-(6-(3-(aminomethyl)phenyl)-1H-indazol-3-yl)-2-cyclopropylacetamide (54 mg, 1 eq, 0.17 mmol) in pyridine (1.5 mL) was added dropwise isobutylsulfonyl chloride (53 mg, 44 μL, 2 eq, 0.34 mmol) at 0 °C. The reaction mixture was stirred at room temperature overnight. Solvent was removed and the residue was partitioned between EtOAc and saturated aqueous sodium bicarbonate solution. The organic layer was dried over $Na_2SO_4$, filtered, and concentrated. The crude residue was purified by preparative HPLC (10−100% MeOH in $H_2O$ + 0.05% TFA) to yield **16** (6 mg, 8% yield) as a colorless solid. $^1H$ NMR (400 MHz, methanol-$d_4$) δ 8.57 (d, $J$ = 0.8 Hz, 1H), 7.84 (dd, $J$ = 8.0, 0.8 Hz, 1H), 7.72 (t, $J$ = 2.0 Hz, 1H), 7.64 – 7.59 (m, 2H), 7.48 (t, $J$ = 7.6 Hz, 1H), 7.42 (d, $J$ = 7.6 Hz, 1H), 4.33 (s, 2H), 2.93 (d, $J$ = 7.2 Hz, 2H), 2.85 (d, $J$ = 6.4 Hz, 2H), 2.16 (dp, $J$ = 13.2, 6.6 Hz, 1H), 1.29 – 1.22 (m, 1H), 1.02 (d, $J$ = 6.8 Hz, 6H), 0.60 – 0.55 (m, 2H), 0.29 (dt, $J$ = 6.0, 4.8 Hz, 2H). $^{13}C$ NMR (214 MHz, Methanol-$d_4$) δ 174.03, 154.48, 144.28, 142.58, 142.00, 140.43, 130.32, 128.63, 128.27, 127.82, 124.53, 121.65, 120.77, 114.96, 61.52, 47.61, 41.01, 26.01, 22.82, 8.00, 4.85. HRMS (ESI): *m/z* calculated for $C_{23}H_{29}N_4O_3S$ $[M+H]^+$: 441.18821. Found: 441.19547.

#### *IN VITRO* STUDIES

##### Kinome Screening.

All compounds except for **16** were profiled via the *scan*MAX assay platform at Eurofins DiscoverX Corporation to assess their selectivity at 1 μM. This selectivity data has been previously reported.[1] The *scan*MAX platform includes 403 wild-type (WT) human kinases and yields percent of control (PoC) values for each compound versus each kinase screened.[2] A selectivity score $S_{10}$(1 μM) can be calculated using these PoC values. Selectivity scores are included in Table 1 and Figure 2. Figure 2 also includes a list of WT human kinases in the *scan*MAX panel with PoC <40 and kinome tree dendrogram generated based upon this selectivity profiling. Data from this *scan*MAX assay platform also enabled generation of Figure 1B, 1C, and 3A.

##### Enzymatic Assays.

Eurofins radiometric enzyme assays were run at the $K_m$ value for ATP for each kinase included in Figure 2C and for CDKL2 in Table 1 to generate the listed $IC_{50}$ values. These assays were run in dose–response (9-pt) format. Assay details for each kinase that was analyzed can be found on the Eurofins website: https://www.eurofinsdiscoveryservices.com.

##### NanoBRET Assays.

Human embryonic kidney (HEK293) cells were acquired from ATCC and maintained in culture using Dulbecco's Modified Eagle's medium (DMEM, Gibco) supplemented with 10% (v/v) fetal bovine serum (FBS, Corning). These cells were incubated at 37°C in 5% $CO_2$, passaging every 72 hours with trypsin (Gibco) to ensure confluency was not reached. Constructs for NanoBRET measurements of CDKL2 (NLuc-CDKL2), AAK1 (NLuc-AAK1), and BMP2K (NLuc-BMP2K) included in Table 1 and Figure 2C were kindly provided by Promega. The N-terminal NLuc orientation of each construct was used in the respective assays. The NanoBRET assays were run in dose–response format as previously described[1] for AAK1 and BMP2K and in accordance with the manufacturer's specifications. Assays were carried out using 0.63 μM of tracer K11 for NLuc-CDKL2, 0.63 μM of tracer K10 for NLuc-AAK1, and 0.13 μM of tracer K10 for NLuc-BMP2K.

Representative normalized curves are included in Figures S1–S3. Where shown, error bars represent standard deviation (Figure S1C and S3).

**Thermal Shift Assays.**
CDKL1, CDKL2, and CDKL3 proteins were prepared using the construct boundaries previously employed to generate their crystal structures (PDB codes: 4AGU, 4AAA, and 3ZDU, respectively).[3] 4 μM of CDKL1, CDKL2, or CDKL3 kinase domain in 10 mM HEPES-NaOH pH 7.4 and 500 mM NaCl was incubated with compound **9** at different concentrations (12.5 or 25 μM) and 5× SyPRO orange dye (Invitrogen). A Real-Time PCR Mx3005p machine (Stratagene) was used to measure fluorescence. A previously reported protocol was followed to run the Tm shift assays and evaluate changes in melting temperatures.[3]

***In Vitro* Kinase Assays.**
For *in vitro* kinase assays, human CDKL1–5 and murine CDKL2 recombinant full-length proteins were used as previously described.[4, 5] Briefly, full-length FLAG-tagged WT plasmids for human CDKL1–5 and mice CDKL2 were obtained from Origene and subsequently subcloned into a pT7CFE1-CHis plasmid (ThermoFisher). The HeLa cell lysate-based Kit (1-Step Human Coupled IVT Kit—DNA, 88881, Life Technologies) was subsequently used for *in vitro* translation and His Pur cobalt spin columns (Thermo Scientific) were used to purify the *in vitro*-translated recombinant protein kinases. SYPRO protein gel staining (ThermoFisher, S12000) and western blot analysis was used to confirm the purity of recombinant proteins. For *in vitro* kinase assays, myelin basic protein (Active Motif, 31314) was employed as a substrate for recombinant kinases. Due to the presence of multiple sites for phosphorylation, myelin basic protein is widely used for *in vitro* kinase assays. Previous studies from others[6, 7] and our groups[5] have shown that CDKL1 and CDKL5 can phosphorylate myelin basic protein *in vitro*. Here, we incubated the purified kinase and myelin basic protein in a kinase buffer (Cell Signaling, 9802) supplemented with or without ATP (50 μM) for 30 min at 30°C, and then measured ADP using the ADP-Glo Kinase Assay kit (Promega). The conversion of ATP to ADP provides a measure of kinase activity in these assays. Under these conditions, experiments were performed using compound **9** in dose–response format (1 nM to 10μM). For data analysis, the kinase activity in the DMSO control group was set as 100% and the relative activity was calculated for compound **9** at various doses.

**Rat Primary Neuron Culture.**
Primary cortical cultures were prepared from embryonic day (E) E17.5 embryos of Sprague Dawley rats, from Charles River. Pregnant females were culled using a $CO_2$ gas chamber followed by cervical dislocation, embryos were removed from the uterus and the brains were taken out. Cortices were dissected out, pooled from multiple embryos, and washed with HBSS. After incubation with 0.25 % trypsin for 15 minutes at 37 °C, trypsin was removed and cortices were blocked with minimum essential medium (MEM) containing 10% fetal bovine serum (FBS), 0.5% dextrose, 0.11 mg/mL sodium pyruvate, 2 mM glutamine and penicillin/streptomycin, for one minute. Cortices were washed once more with HBSS. Cells were dissociated and then counted using a hemocytometer. Neurons were plated on 12-well culture plates at a density of 200,000 cells per well. The wells were coated with 50mg/ml PLL and placed in the incubator overnight. Neurons were plated in neurobasal medium containing 1 mL of B27 (Gibco), 0.5 mM glutamax, 0.5 mM glutamine, 12.5 μM glutamate and penicillin/streptomycin. Primary neuronal cultures were kept at 37 °C and 5% $CO_2$. At DIV15, neurons were treated with 0.5 μM, 1.0 μM, 2.0 μM

and 4.0 μM of the compound **9** ± 50 nM of the SGC-CAF382-1 compound or with 1 μM and 10 μM of compound **16** compound for 1 hour. The compounds were added directly to the media and the plates were place at 37 °C for the time of the treatment. DMSO was added to the well for the control condition.

**Rat Primary Neuron Western Blot.**
After treatment, neuronal cultures were lysed in 300 μL of 2X sample buffer (Invitrogen) containing 0.1 M DTT. Lysates were twice sonicated briefly and denatured at 70 °C for 10 minutes. The samples were centrifuged at 13,300 rpm for 10 minutes and ran on NuPage 4-12% Bis-Tris polyacrylamide gels (Invitrogen). Proteins were transferred onto a Immobilon PVDF membrane (Millipore), which was then blocked in 5% milk in tris-buffered saline containing 0.1% Tween-20 (TBST) at 4 °C overnight. Primary antibodies were incubated at room temperature for 90 minutes, and HRP-conjugated secondary antibodies at RT for 2 hours. The following primary antibodies were used: rabbit anti-pS222 EB2 (1:1,000; Covalab, from [8]), rat anti-EB2 (1:2,000; Abcam ab45767), mouse anti-tubulin (1:10,000; Sigma T9026). The following secondary antibodies were used at a concentration of 1:10,000: HRP-conjugated anti-rabbit (Jackson 711-035-152), HRP-conjugated anti-mouse (Jackson 715-035-151) and HRP-conjugated anti-rat (Jackson 712-035-153). The membrane was developed using ECL reagent (Cytiva) and was visualized with an Amersham Imager 600 (GE Healthcare). Quantification of Western blots in Figures 5 and S4 was manually performed using ImageJ Software (version 1.54d). EB2 phosphorylation was measured relative to total EB2.

**Mammalian Cell Culture Conditions.**
MDA-MB-231 cells (ATCC) were cultured in DMEM (high-glucose) containing 10% fetal bovine serum and MCF-7 cells (ATCC) were cultured in Alpha MEM containing 10% fetal bovine serum, 1 mM Na/Pr, 10 μg/ml insulin, and 1% NEAA.

TU-BcX-4IC cells were cultured in RPMI 1640 medium (Gibco, 72400-047) containing 10% fetal bovine serum (Gibco, 10437-028), 1% MEM amino acids solution (Gibco, 11130-051), 1% MEM non-essential amino acids (Sigma-Aldrich, M1745), and 1% antibiotic-antimycotic (Gibco, 15240-062).

**Cell Viability.**
MDA-MB-231 and MCF-7 cells were plated at a seeding density of 1,200 cells per well on a 384 well plate and were incubated overnight to adhere. Compounds were added to these cells in quadruplicate with 0.1% DMSO as the negative control and 10% DMSO as the positive control. TU-BcX-4IC cells were seeded at a density of 2,500 cells per well on a half-area 96-well plate and were incubated overnight to adhere. The following day, compounds were added to adhered TU-BcX-4IC cells in duplicate with 0.1% DMSO as the negative control and 10% DMSO as the positive control. After 48 hours, CellTiter-Glo reagent (Promega) was added to each well. Plates were read on a GloMax Discover (Promega) or BioTek Synergy. Percent viability was calculated using the following formula: ((raw RFU value – average of positive control wells)/(average of negative control wells – average of positive control wells))*100. Plots and/or $IC_{50}$ estimates in Figure S6 were generated using GraphPad Prism.

**Crystal Violet Staining.**

TU-BcX-4IC cells were seeded at a density of 9,000 cells per well on a 48-well plate (8,000 cells/cm$^2$) and were incubated overnight to adhere. The following day, compounds were added in duplicate with DMSO as the negative controls at each log dose concentration. After 48 hours, cells were washed once with 1X sterile PBS. Cells were then stained with 3% crystal violet (Sigma C0775) in 100% methanol (Spectrum 67-56-1). After 30 minutes, the stain was removed, and cells were washed with DI water until clear. After drying overnight, cells were imaged at 4x magnification using an Agilent Cytation 10 Confocal Imaging System (Santa Clara, California). Images were quantified using ImageJ software (National Institutes of Health, Bethesda, Maryland, USA, http://imagej.nih.gov/ij/). Total cell area and average cell size values (Figure S6) were normalized to DMSO control values at corresponding concentrations and plotted using GraphPad Prism.

**Mammalian Cell Western Blot.**

Cells were plated on 6-well plates with a seeding density of 175,000 and 200,000 cells per well for MDA-MB0231 and MCF-7 cells, respectively. Compounds were added after allowing the cells to adhere overnight. After 48 hours, cells were lysed in RIPA buffer containing 1X protease inhibitor cocktail and sonicated. Blots were blocked in 5% milk in 1X TBST (0.1% Tween20) for 1 hour at room temperature. Primary antibodies were diluted in 5% BSA in 1X TBST and incubated overnight: 1:5000 Actin (Sigma 2228); 1:5000 CD44 (Proteintech 15675-1-AP); 1:500 ZEB1 (Proteintech 21544-1-AP); 1:1000 β-catenin (Proteintech 51067-2-AP). Secondary antibodies were diluted in 5% milk in 1X TBST and incubated at room temperature for 1 hour. Blots were imaged on the iBright (ThermoFisher Scientific). Blots in Figure S5 were analyzed using Fiji ImageJ. Band densities were normalized to Actin housekeeping control. Plots were generated using GraphPad Prism.

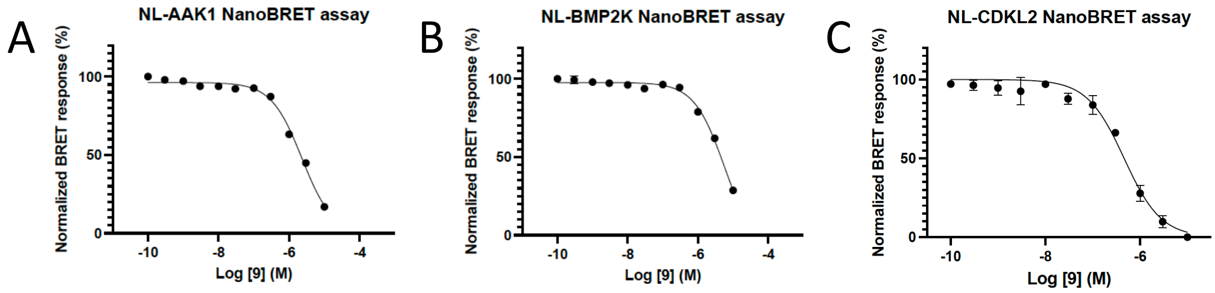


**Figure S1.** AAK1, BMP2K, and CDKL2 NanoBRET assay curves corresponding with the data in Table 1 and Figure 2 for compound **9**. AAK1 and BMP2K NanoBRET assays were run in singlicate (n = 1), while the CDKL2 NanoBRET assay was run in triplicate (n = 3). Error bars represent standard deviation (SD).

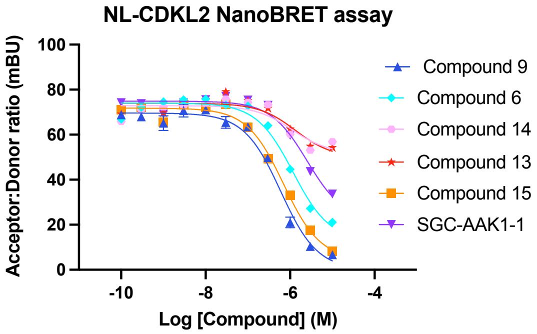


**Figure S2.** CDKL2 NanoBRET assay curves for compounds in Table 1 (n = 1).

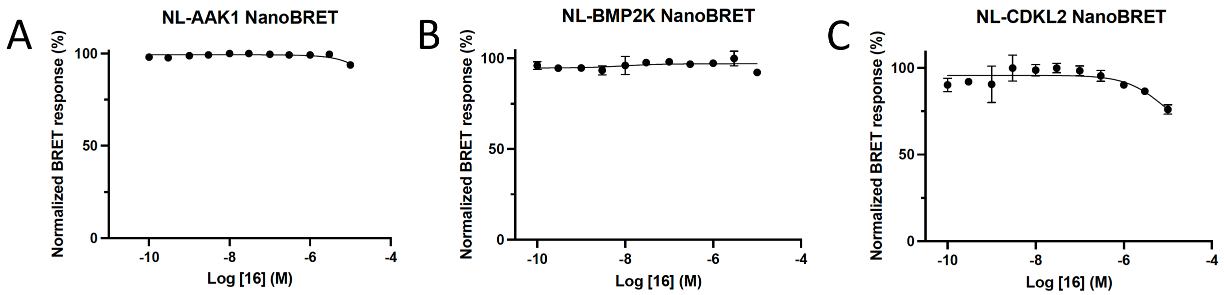


**Figure S3.** AAK1, BMP2K, and CDKL2 NanoBRET assay curves corresponding with the data in Table 1 for compound **16**. All assays were run in singlicate (n = 1), Error bars represent standard deviation (SD).

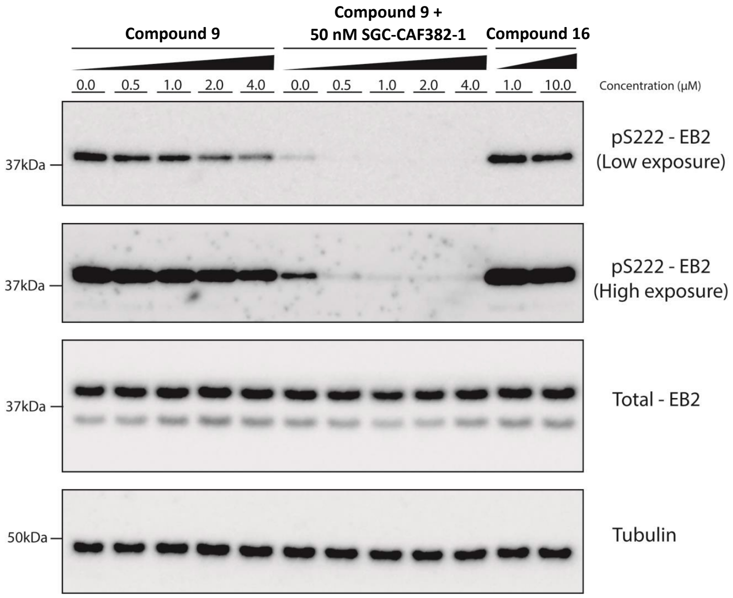


**Figure S4.** Second replicate for Western blot analysis of pS222-EB2 in primary rat neurons after 1 h treatment with compound **9**, SGC-CAF382-1, compound **9** + SGC-CAF382-1, or compound **16**.

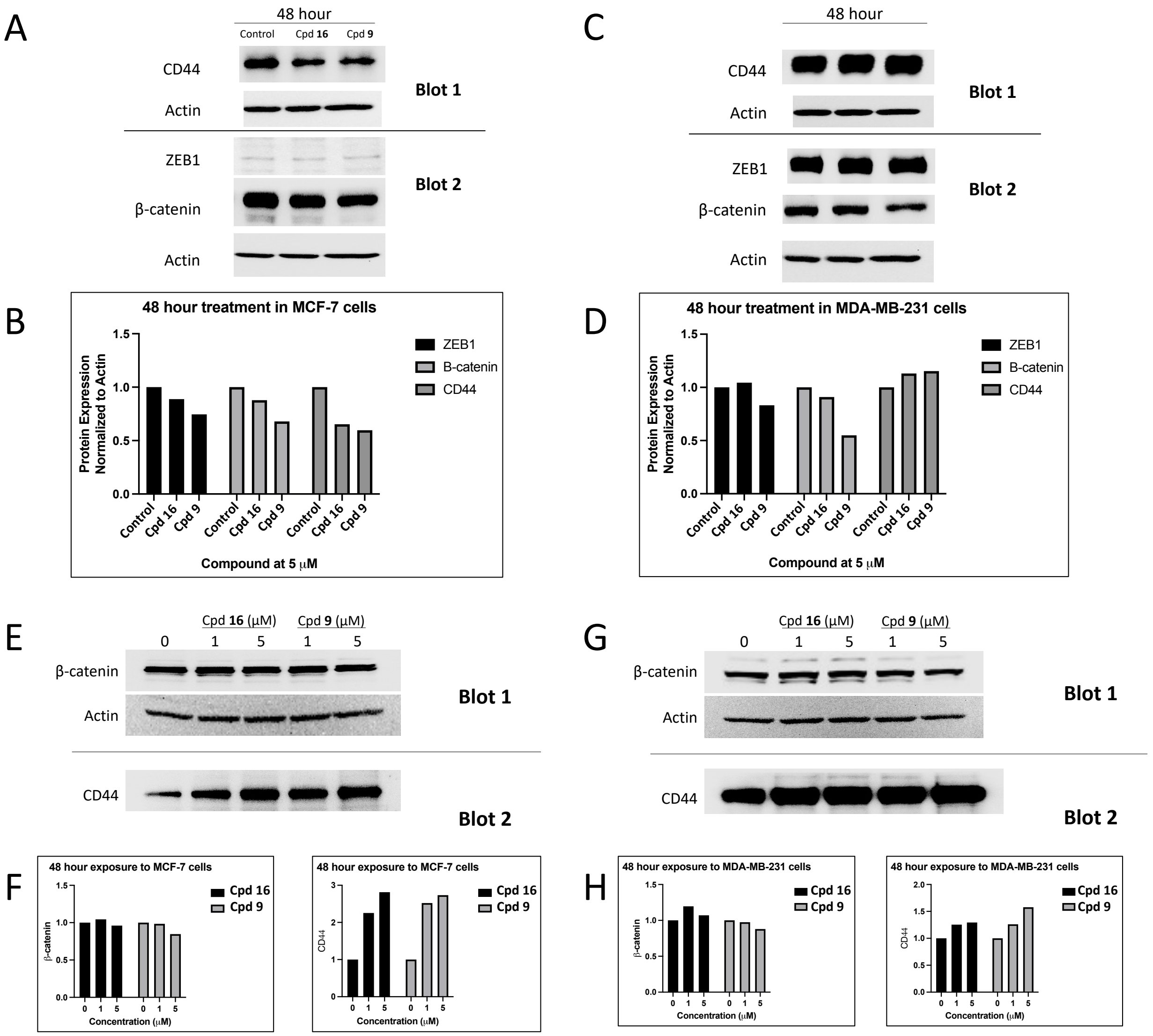


**Figure S5.** Example Western blots generated to probe the response of EMT proteins to treatment of breast cancer cell lines with compounds **9** and **16** for 48 hours. (A) Western blots showing the expression of CD44, actin, ZEB1, or β-catenin in response to control (DMSO), 5 μM compound **9**, or 5 μM compound **16** after 48h treatment in MCF7 cells. (B) Quantification of the Western blots shown in panel A. (A) Western blots showing the expression of CD44, actin, ZEB1, or β-catenin in response to control (DMSO), 5 μM compound **9**, or 5 μM compound **16** after 48h treatment in MDA-MB-231 cells. (D) Quantification of the Western blots shown in panel C. (E) Western blots showing the expression of CD44, actin, ZEB1, or β-catenin in response to control (DMSO), 5 μM compound **9**, or 5 μM compound **16** after 48h treatment in MCF7 cells. (E) Western blots showing the expression of CD44, actin, ZEB1, or β-catenin in response to control (DMSO), 1 or 5 μM compound **9**, or 1 or 5 μM compound **16** after 48h treatment in MCF7 cells. (F) Quantification of the Western blots shown in panel E. ZEB1 was excluded from analysis. (G) Western blots showing the expression of CD44, actin, ZEB1, or β-catenin in response to control (DMSO), 1 or 5 μM compound **9**, or 1 or 5 μM compound **16** after 48h treatment in MDA-MB-231 cells. (H) Quantification of the Western blots shown in panel G. ZEB1 was excluded from analysis.

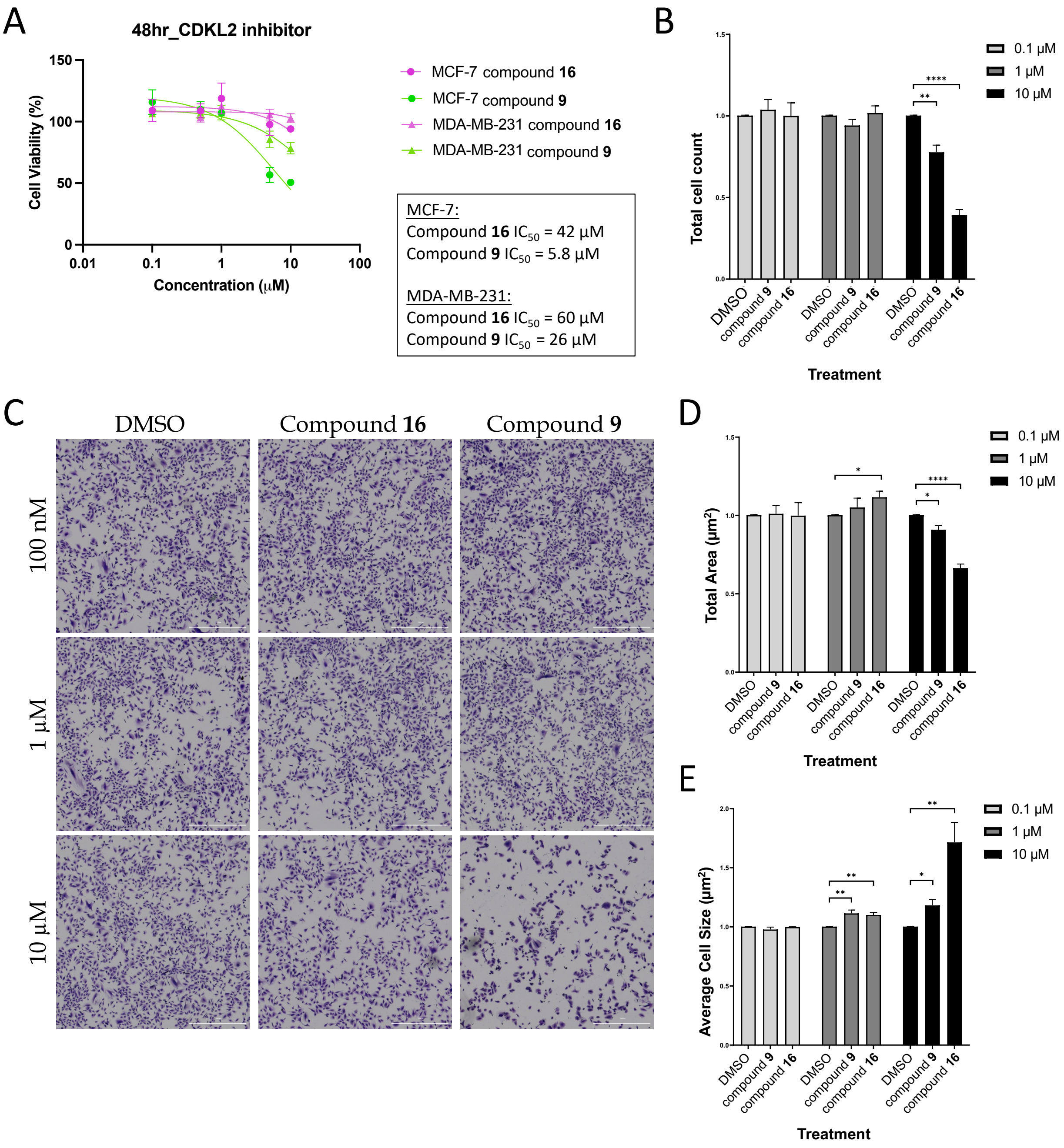


**Figure S6.** Viability data for CDKL2 probe and negative control in breast cancer cell lines. (A) Viability curves and corresponding $IC_{50}$ values generated when MCF7 and MDA-MB-231 cells were treated in dose–response format for 48 hours with either compound **9** or **16**. Compound **9** was only cytotoxic in MCF7 cells up to 10 µM with an absolute IC50 = 5.8 µM. (B) Viability data generated when TU-BcX-4IC cells were treated in dose–response format for 48 hours with either compound **9** or **16**, n = 4. (C) Representative images at 4X magnification of crystal violet stained TU-BcX-4IC cells treated with either compound **9** or **16** in dose–response format for 48 hours. (D) Quantification of total cell area when TU-BcX-4IC cells were treated in dose–response format for 48 hours with either compound **9** or **16**, n = 4. (E) Quantification of average cell size when TU-BcX-4IC cells were treated in dose–response format for 48 hours with either compound **9** or **16**, n = 4. For panels (D) and (E), values are plotted as the mean ± SEM normalized to the DMSO-treated control, * $p < 0.05$; ** $p < 0.01$; *** $p < 0.001$; **** $p < 0.0001$.

#### CRYSTALLOGRAPHY METHODS

**Table S1.** Statistics for CDKL2 data collection, phasing, and refinement.

| | |
|---|---|
| **Data Collection Statistics** | |
| Radiation source | Diamond I04 |
| Wavelength (Å) | 0.9537 |
| Spacegroup | P $3_2$ |
| Cell dimensions: | |
| *a*, *b*, *c* (Å) | 58.09 58.09 104.26 |
| *α*, *β*, *γ* (°) | 90 90 120 |
| Number of molecules/asymmetric unit | 1 |
| Resolution range (Å) | 52.13-1.72 (1.75-1.72) |
| Total observations | 818760 (20848) |
| Unique reflections | 41802 (2106) |
| Completeness (%) | 100 (100) |
| Multiplicity | 19.6 (9.9) |
| $R_{merge}^a$ | 0.059 (2.619) |
| Average I/σ *(I)* | 22.7 (0.4) |
| $CC_{1/2}$ (%) | 99.9 (50.8) |
| **Refinement and model statistics** | |
| Resolution range (Å) | 50.310 - 1.721 (1.782 - 1.721) |
| Number of reflections used | 41694 (4110) |
| $R_{work}^b/R_{free}^c$ (%) | 20.46/22.19 |
| Ligand ID | LIG |
| **B values ($Å^2$)** | |
| Overall | 49.67 |
| Macromolecules | 49.62 |
| Ligands | 48.98 |
| Waters | 50.99 |
| **Root mean square deviation from ideality** | |
| Bond lengths (Å) | 0.005 |
| Bond angles (°) | 0.832 |
| **Number of atoms** | |
| Protein atoms | 2465.00 |
| Ligands | 43.00 |
| Waters | 119.00 |
| **Ramachandran analysis** | |
| Favored regions / Allowed regions / Outliers (% of residues) | 97.65/2.35/0.00 |

| | |
|---|---|
| Rotamer outliers (%) | 0.37 |
| Clashscore | 4.20 |
| **PDB Code** | 8S6I |

*Values in parentheses are for the highest resolution shell.

**CDKL2 Protein Expression and Purification.**

Baculoviral expression of human CDKL2 (UniProt Q92772; residues 1–308; T159D and Y161E) was performed in Sf9 cells as described previously.[3] Cells were harvested 72 hr post-infection and resuspended in 30 mL binding buffer (50 mM HEPES pH 7.5, 500 mM NaCl, 5% glycerol, and 1 mM TCEP) supplemented with protease inhibitors. Cells were lysed via sonication. Benzonase (1:1000) was added to cleave DNA. The cell lysate was clarified by centrifugation and proteins purified by nickel-affinity, size exclusion, and ion exchange chromatography. The CDKL2 buffer was exchanged to size exclusion buffer (10 mM HEPES, pH 7.5, 500 mM NaCl, 5% glycerol, and 1 mM TCEP) by using a PD10 column. CDKL2 was then diluted to 1 mg/mL, followed by addition of compound **9** in 3-fold molar excess with 3 hours incubation at 4°C. CDKL2 was concentrated to 10 mg/mL for crystallization studies.

**Crystallization.**

Crystals were grown in 0.09 M sodium fluoride, 0.09 M sodium bromide, 0.09 M sodium iodide, 0.1 M Tris, 0.1 M BICINE, pH 8.5, 33% (v/v) ethylene glycol and 16% (w/v) PEG 8000 at 4°C, appearing after 4 days. 25% ethylene glycol was added as a cryoprotectant for crystal mounting before vitrification in liquid nitrogen.

**Diffraction Data Collection, Structure Solution and Refinement.**

Diffraction data were collected on beamline i04 at Diamond Light Source to a resolution of 1.72 Å. Data were processed with the Xia2 pipeline[9], using DIALS software.[10] PDB 4AAA was employed as a search model for molecular replacement in PHENIX.[11] The initial model was improved through iterative rounds of manual building in Coot[12] and refined by PHENIX.[11] Data collection and refinement statistics are included in Table S1. Interactions were mapped using Protein–Ligand Interaction Profiler (PLIP).[13]

[1]H NMR of *N-(6-bromo-1H-indazol-3-yl)cyclopropanecarboxamide* (**17**) in DMSO-$d_6$

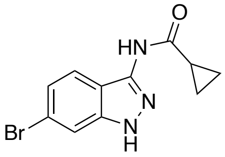

**17**

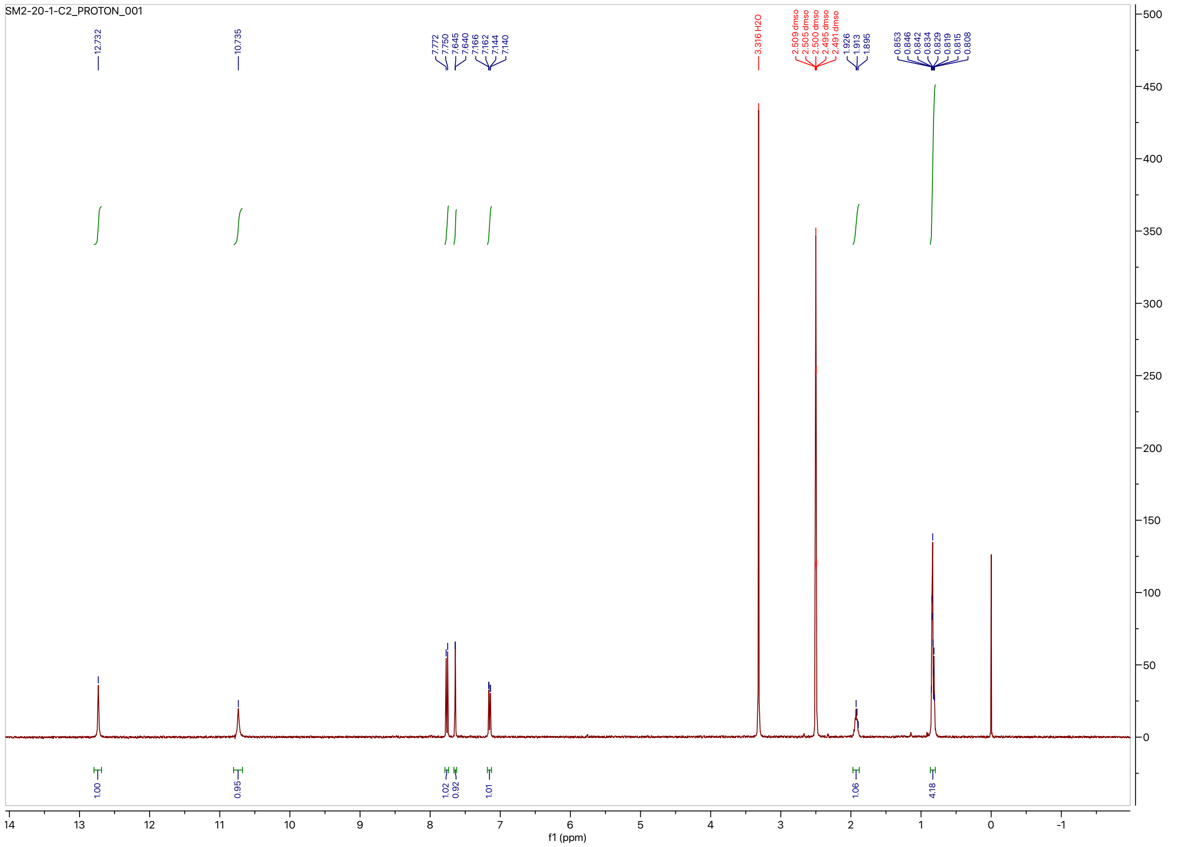

$^{13}$C NMR of *N-(6-bromo-1H-indazol-3-yl)cyclopropanecarboxamide* (**17**) in DMSO-$d_6$

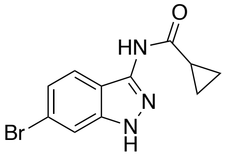


**17**

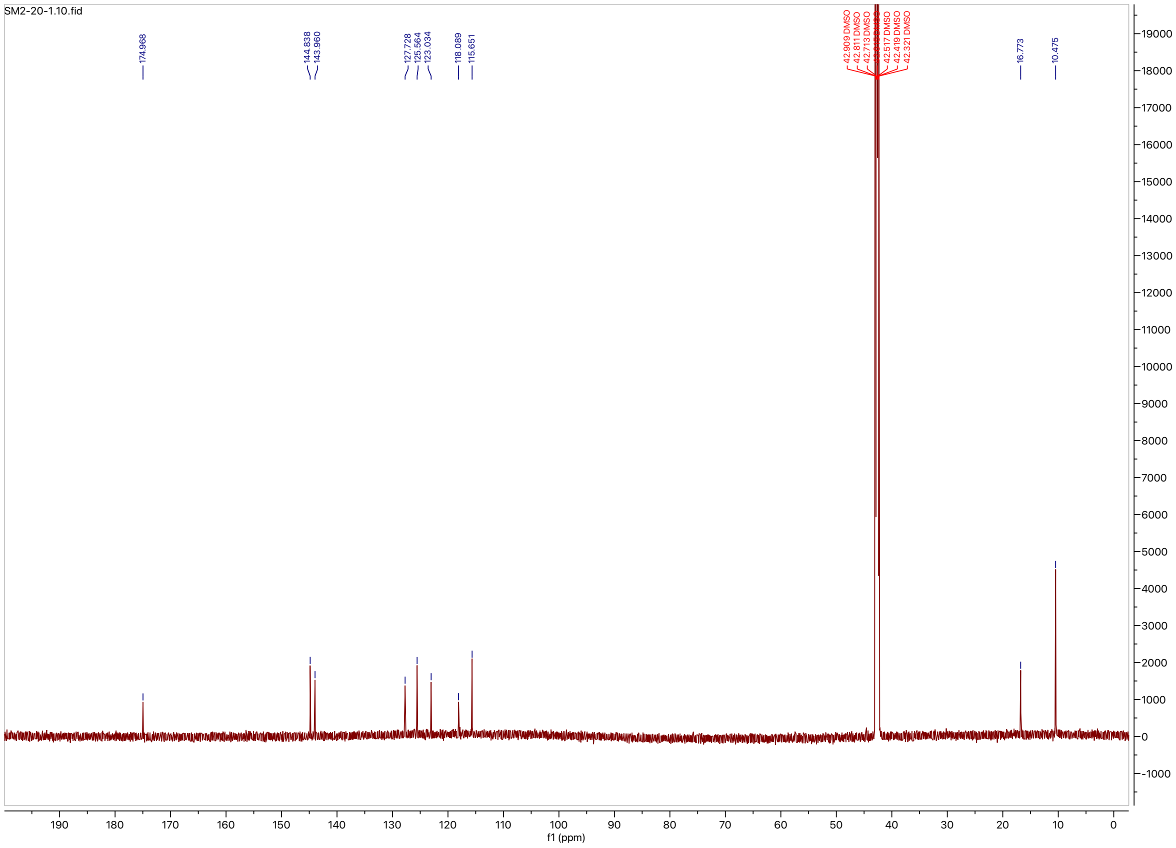

[1]H NMR of *N*-(6-(3-(((2-methylpropyl)sulfonamido)methyl)phenyl)-1*H*-indazol-3-yl)cyclopropanecarboxamide (**9**) in DMSO-$d_6$

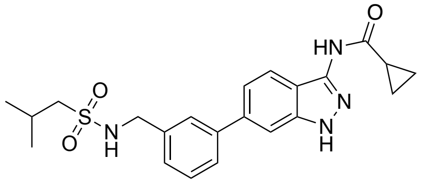

**9**

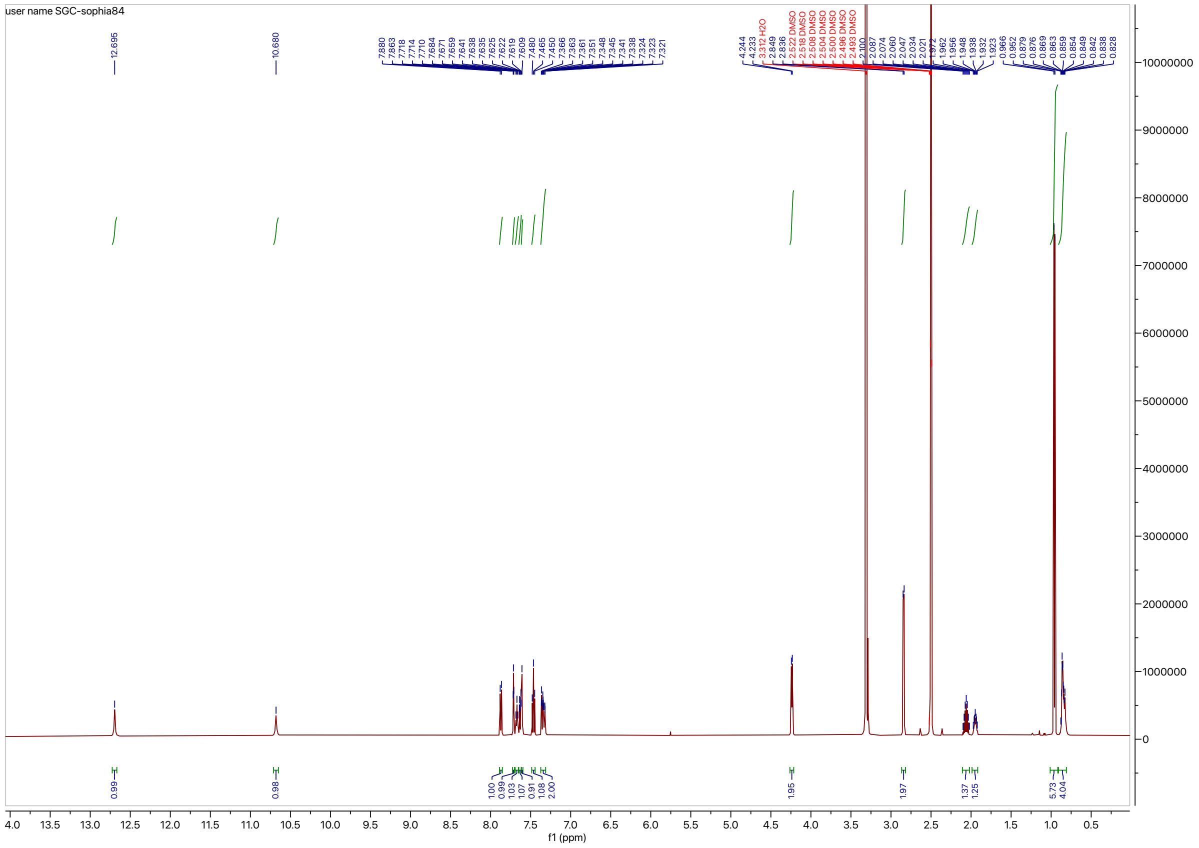

$^{13}$C NMR of *N*-(6-(3-(((2-methylpropyl)sulfonamido)methyl)phenyl)-1*H*-indazol-3-yl)cyclopropanecarboxamide (**9**) in DMSO-$d_6$

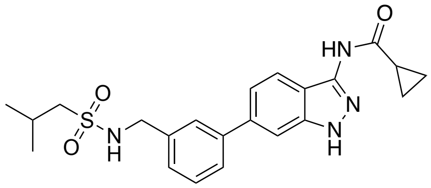

**9**

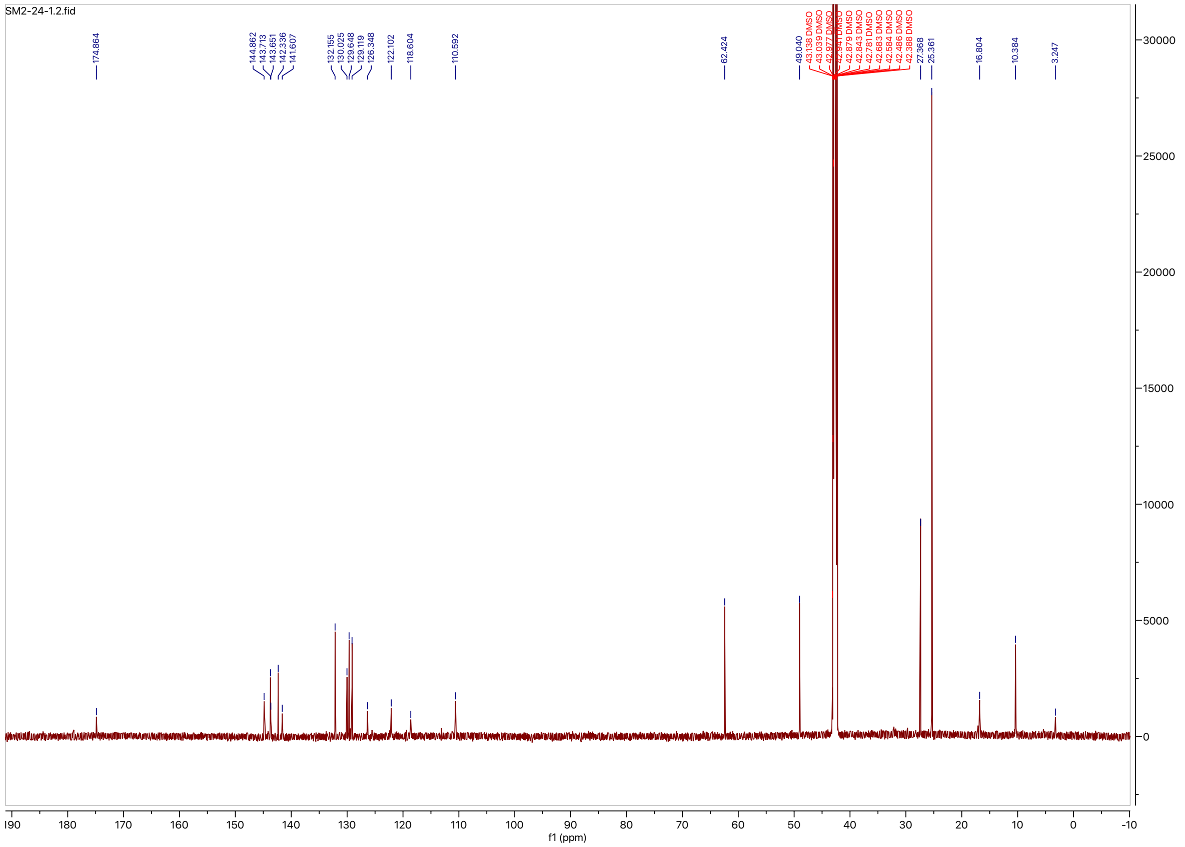

$^{1}$H NMR of *N*-(6-bromo-1*H*-indazol-3-yl)-2-cyclopropylacetamide (**18**) in methanol-$d_4$

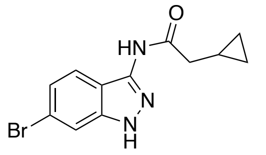


**18**

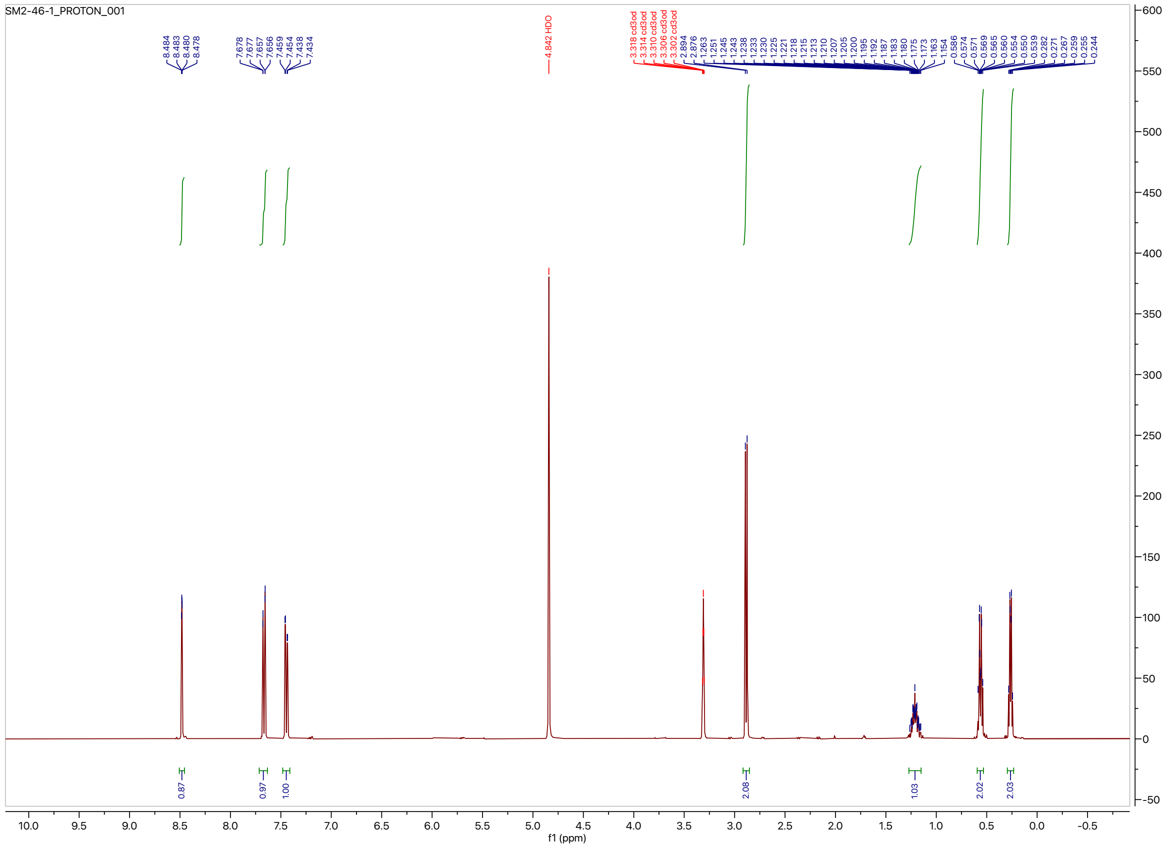

$^{13}$C NMR of *N*-(6-bromo-1*H*-indazol-3-yl)-2-cyclopropylacetamide (**18**) in methanol-$d_4$

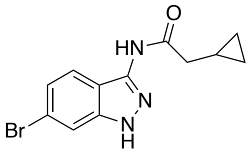

**18**

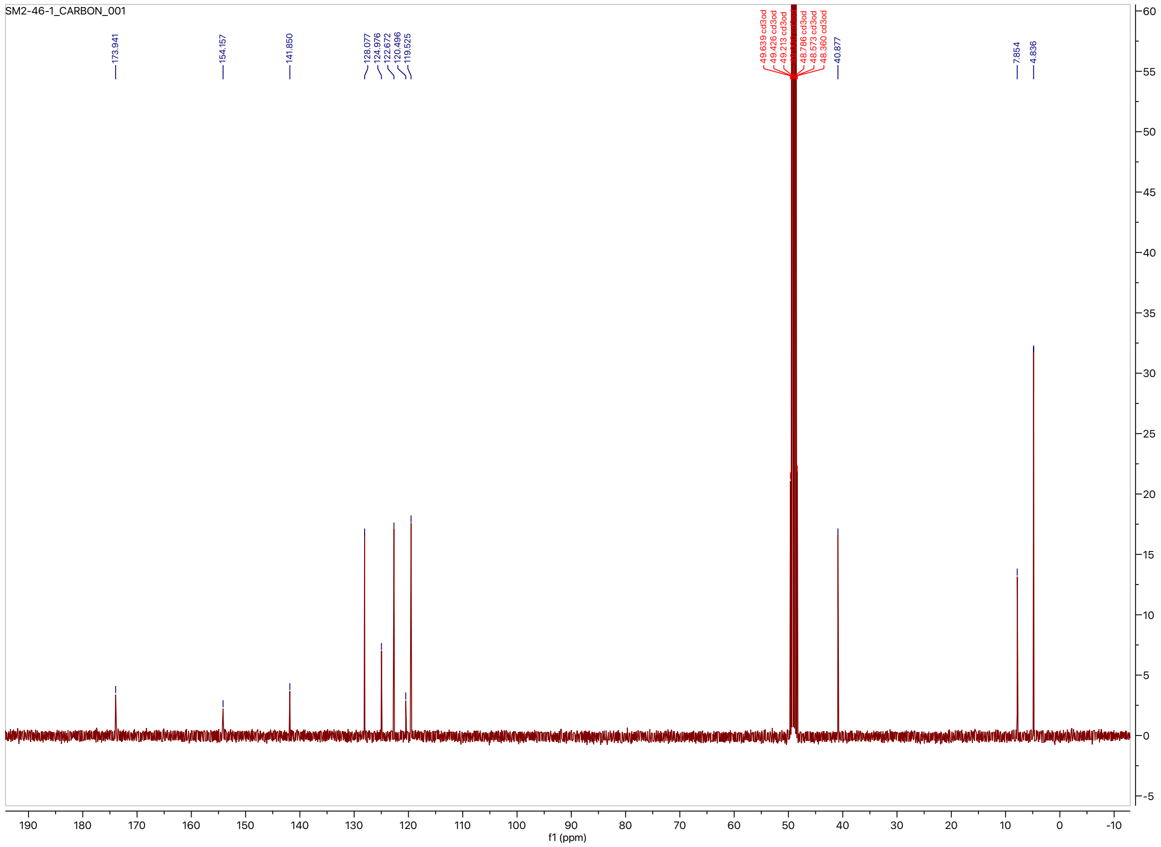

[1]H NMR of 2-cyclopropyl-*N*-(6-(3-(((2-methylpropyl)sulfonamido)methyl)phenyl)-1*H*-indazol-3-yl)acetamide (**16**) in methanol-$d_4$

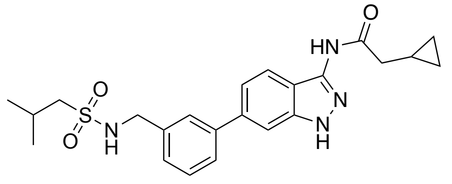

**16**

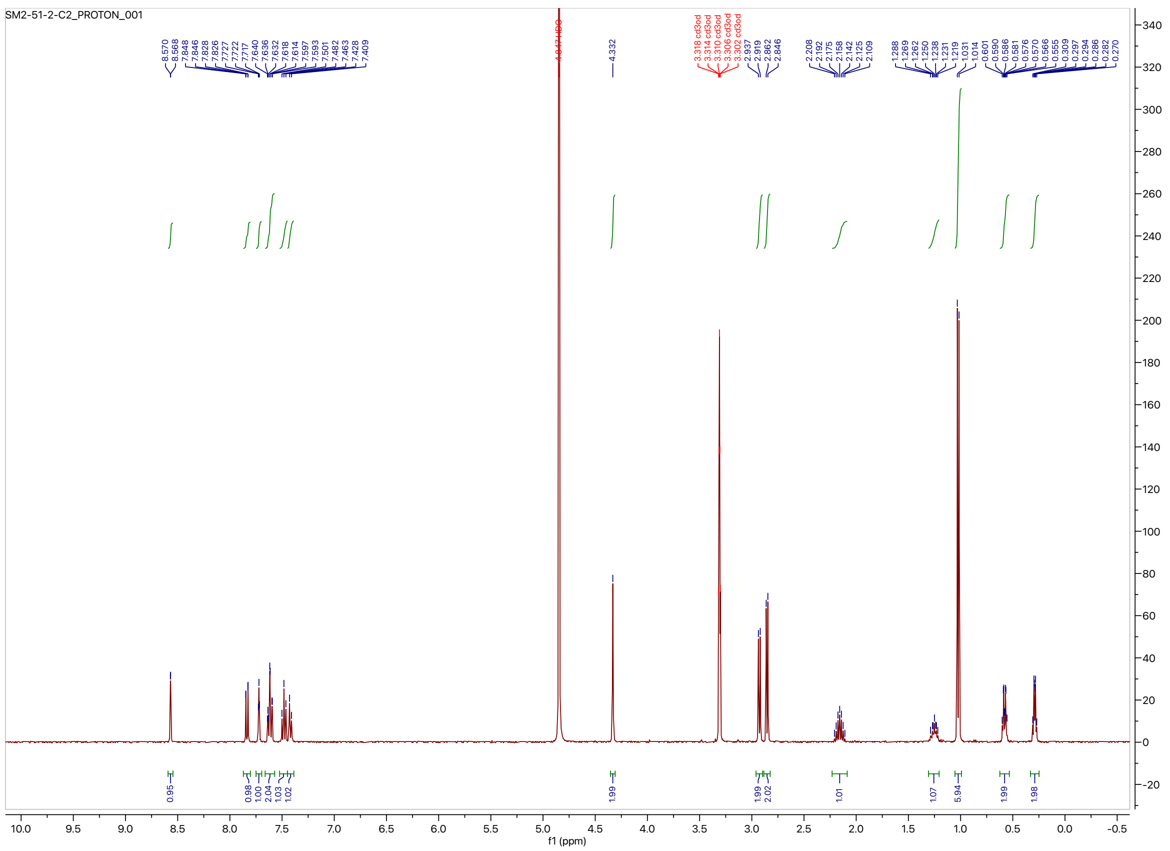

$^{13}C$ NMR of 2-cyclopropyl-*N*-(6-(3-(((2-methylpropyl)sulfonamido)methyl)phenyl)-1*H*-indazol-3-yl)acetamide (**16**) in methanol-$d_4$

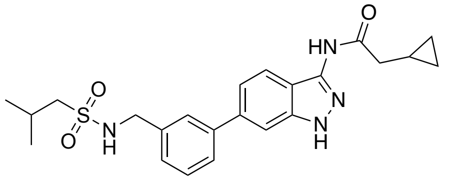

**16**

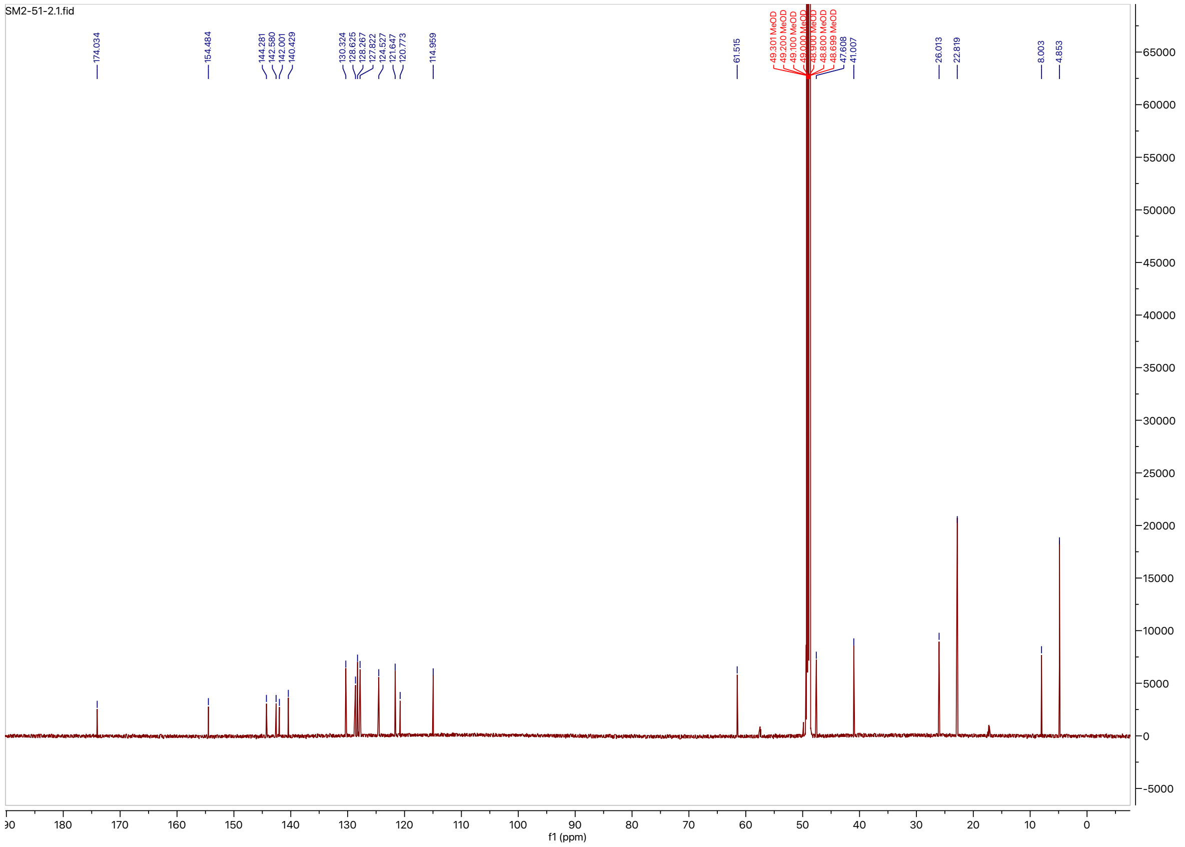
